# Regulation and dynamics of force transmission at individual cell-matrix adhesion bonds

**DOI:** 10.1101/530469

**Authors:** Steven J. Tan, Alice C. Chang, Cayla M. Miller, Sarah M. Anderson, Louis S. Prahl, David J. Odde, Alexander R. Dunn

**Affiliations:** Department of Chemical Engineering, Stanford University, Stanford, CA 94305; Department of Biomedical Engineering and Physical Sciences-Oncology Center, University of Minnesota, Minneapolis, MN 55455

## Abstract

Integrin-based adhesion complexes link the cytoskeleton to the extracellular matrix (ECM) and are central to the construction of multicellular animal tissues. How biological function emerges from the 10s-1000s of proteins present within a single adhesion complex has remained unclear. We used fluorescent molecular tension sensors to visualize force transmission by individual integrins in living cells. These measurements revealed an underlying functional modularity in which integrin class controlled adhesion size and ECM ligand specificity, while the number and type of connections between integrins and F-actin determined the force per individual integrin. In addition, we found that most integrins existed in a state of near-mechanical equilibrium, a result not predicted by existing models of cytoskeletal force transduction. A revised model that includes reversible crosslinks within the F-actin network accounts for this result, and suggests how cellular mechanical homeostasis can arise at the molecular level.

## Main Text

Integrins are heterodimeric transmembrane proteins that form the core of micron-sized protein assemblies, here referred to generically as focal adhesions (FAs), that link the cytoskeleton to the extracellular matrix (ECM) and hence play a central role in the construction of multicellular tissues *(1–4)*. Proteomics studies demonstrate that ~60 proteins constitute the core integrin adhesion machinery, and that >2,400 proteins are potential members of the integrin adhesome *(5)*. Previous studies have uncovered a dense web of interactions between FA proteins *(6)*, the complexity of which poses a challenge in understanding how FAs function as an integrated whole.

In this study, we sought to better understand how FA-mediated force transmission arises at the molecular level. The transmission of forces between the cytoskeleton and ECM constitutes a core function of FAs and is required both for tissue morphogenesis and many forms of cell migration. Force transmission is commonly described in terms of the molecular clutch model, in which continuous slippage between the rearward flowing actin cytoskeleton and FA components mediates force transmission to the ECM *(7–10)*. This model reproduces important biological observations, for example biphasic traction forces as a function of substrate stiffness *(11–13)*. However, to our knowledge the clutch model has not been directly tested by the observation of the dynamics of force transmission at the single-molecule level in living cells.

To address this gap, we used FRET-based molecular tension sensors (MTSs) to measure the loads experienced by individual integrin heterodimers in a variety of cell types (Fig. 1a-d). MTS_low_ and MTS_FN9-10_ report on loads between 2 and 7 pN and present either a linear RGD peptide or the fibronectin type III domains 9 and 10, respectively *(14, 15)*. In addition, we developed a new sensor termed MTS_high_ that measures forces between 7 and 11 pN and contains the same RGD motif as MTS_low_ (Fig. S1-S2) *(16)*.

**Fig. 1.**
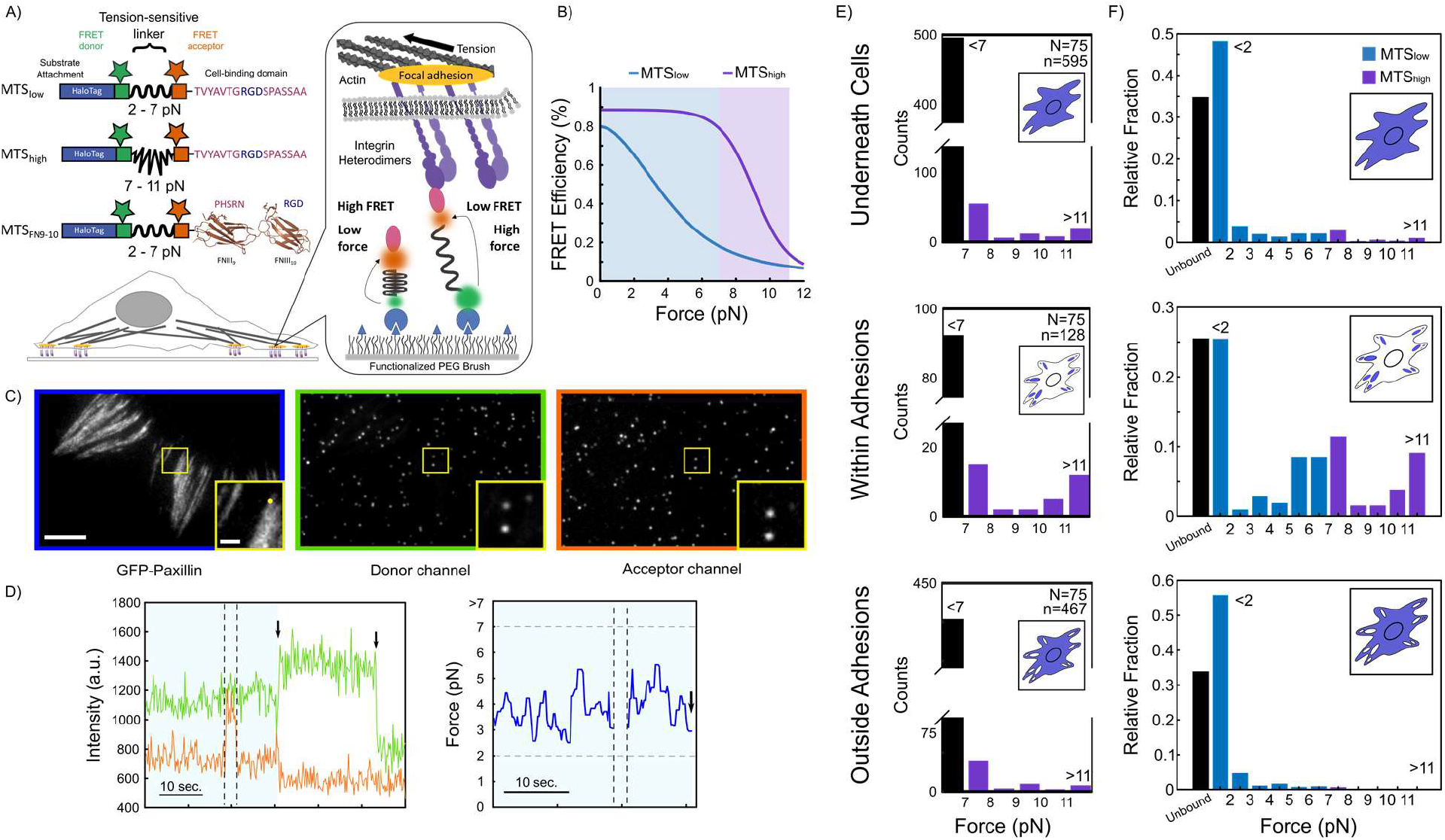
Single molecule tension measurements in living cells reveal distinct subpopulations of load-bearing integrins. (A) FRET-based molecule tension sensors (MTSs). MTSs are attached to the coverslip surface via the HaloTag domain. (B) FRET-force calibration curves for MTS_low_ (blue) and MTS_high_ (purple) *(16, 42)*. (C) Representative images showing GFP (left), FRET donor (middle), and acceptor channels (right) for HFFs adhering to a surface functionalized with MTS_low_. Scale bar = 5 μm, inset scale bar = 2 μm. (D) Example intensity traces (left) for the FRET donor (green) and acceptor (orange). Dashed lines delineate frames during which the acceptor dye was directly excited with 633 nm light; arrows mark acceptor or donor bleaching. Right: corresponding load timeseries prior to acceptor photobleaching (light blue). (E) Single molecule load distributions for MTS_high_ underneath cells, within adhesions, and outside adhesions. *N* = number of cells, *n* = number of sensors (F) Combined single-molecule load distributions for MTS_low_ and MTS_high_ sensors underneath cells, within adhesions, and outside adhesions for MTS_low_ (blue; data from ref. *(14)*) and MTS_high_ (purple).

In previous studies, we found that a majority of ligand-bound integrins exist in a minimally tensioned state that does not depend on the actin cytoskeleton *(14)*, which we confirm in this study (Fig. 1e). Measurements using MTS_high_ revealed that the distribution of loads on individual integrins was highly asymmetric, with a small minority of integrins within the adhesions of human foreskin fibroblasts (HFFs) bearing loads of ~6 pN and >10 pN (Fig. 1e, 1f). The presence of the latter subpopulation is consistent with previous studies demonstrating that at least some integrins experience peak loads >50 pN *(17–21)*.

How these different load subpopulations arose at the molecular level was unclear. One plausible explanation was that these subpopulations might correspond to ligation by different integrin heterodimers, a scenario supported by studies reporting distinct roles for α_5_β_1_ and α_v_β_3_ integrins in adhesion and traction generation *(22–25)*. To test this hypothesis, we made use of pan-integrin knockout (pKO) mouse embryonic fibroblasts (MEFs) rescued with the integrin α_v_ subunit (pKO-α_v_), which forms predominantly α_v_β_3_ and α_v_β_5_ heterodimers, the β_1_ subunit (pKO-β_1_), which forms only α_5_β_1_ integrin in these cells, or both subunits (pKO-α_v_/β_1_), which form all three integrin heterodimers *(22)*. pKO-α_v_ and pKO-α_v_/β_1_ cells spread normally on coverslips functionalized with either MTS_low_ or MTS_high_ and formed sizeable FAs (Fig. 2a, top and middle, Fig. S3, S4) while the majority of pKO-β_1_ cells failed to spread on either sensor (see insets). In contrast, all three cell types spread on coverslips functionalized with MTS_FN9-10_, though pKO-α_v_ cells yielded lower integrated traction forces, without significant changes in adhesion size (Fig. 2a, bottom, Fig. S4, S5). Thus, integrin usage and ligand identity strongly influenced adhesion and traction generation at the whole-cell level, an outcome consistent with previous observations.

**Fig. 2.**
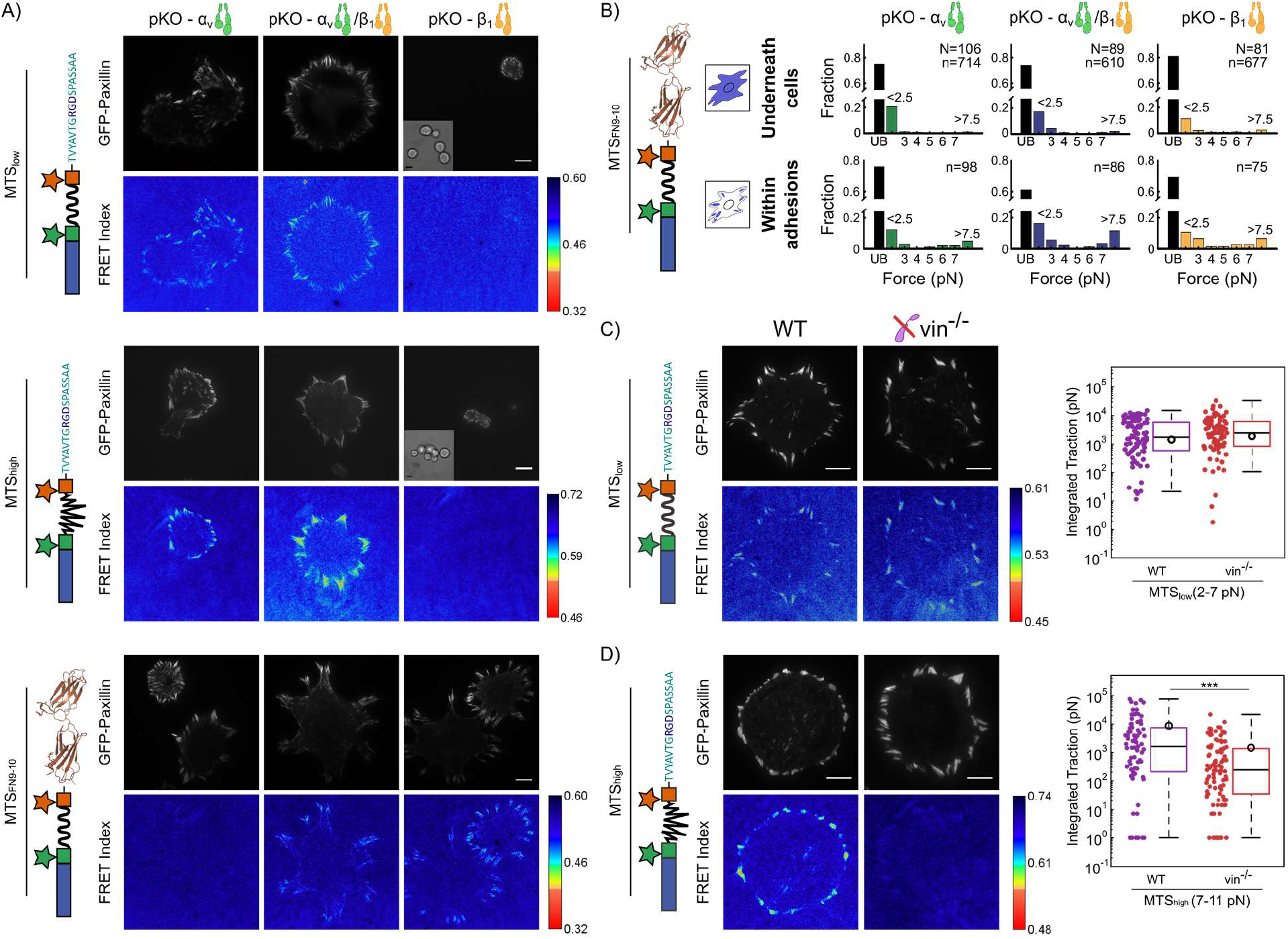
Force per integrin is independent of integrin class, but is influenced by linkages to F-actin. (A) Images of GFP-paxillin and ensemble FRET maps for pKO-α_v_, pKO-β_v_/β_1_, and pKO-β_1_ cells adhering to MTS_low_ (top), MTS_high_ (middle), and MTS_FN9-10_ (bottom). Insets show corresponding brightfield images for pKO-β_1_ cells, which rarely spread on surfaces functionalized with MTS_low_ and MTS_high_. Scale bars = 10 μm. (B) Single-molecule load distributions for pKO cell lines adhering to MTS_FN9-10_. Black bars indicate unbound molecules. *N* = number of cells, *n* = number of sensors. (C, D, left) Ensemble FRET maps for WT and vin^−/−^ MEFs transfected with GFP-paxillin and seeded on coverslips functionalized with MTS_low_ and MTS_high_ sensors. Scale bar is 10 μm. (C, D, right) Total integrated traction per cell for forces <7 pN measured with MTS_low_. Open circles indicate the mean value. (WT: 96 cells, mean: 3.6 nN; vin^−/−^: 89 cells, mean: 4.4 nN.) Total integrated traction per cell for forces between 7 and 11 pN measured with MTS_high_. Open circles indicate the mean. (WT: 71 cells, mean: 8.7 nN; vin^−/−^: 99 cells, mean: 1.5 nN.) (\*\*\**p* < 0.001 using two-sided Wilcoxon ranksum test.)

We next measured the distribution of loads experienced by individual integrins for pKO-α_v_, pKO-β_1_, and pKO-α_v_/β_1_ cells adhering to coverslips functionalized with MTS_FN9-10_. Contrary to expectation, the distributions of loads for integrins bearing >_2_ pN were strikingly similar across all three cell lines (Fig. 2b, Fig. S6). However, the fraction of integrin-bound sensors underneath cells was significantly higher for pKO-α_v_/β_1_ and pKO-β_1_ cells as compared to pKO-α_v_ cells (Fig. 2b, Table S1), a factor that can largely account for the differences in traction generation at the whole-cell level. Thus, integrin usage indirectly influenced overall cellular traction by modulating the fraction of engaged integrins, but did not influence the distribution of loads borne by individual, ligand-bound integrins.

An alternate hypothesis was that the load experienced by an integrin, regardless of class, is determined by the nature of its linkages to the actin cytoskeleton. The cytosolic protein vinculin reinforces the talin-actin linkage and may influence the load borne by individual integrins. We quantified traction generation at the whole-cell and single-integrin level for wild-type (WT) and vinculin-null (vin^−/−^) MEFs adhering to MTS_low_ and MTS_high_ (Fig. 2c, 2d). Importantly, WT but not vin^−/−^ MEFs generated appreciable regions with low FRET when adhering to MTS_high_ (Fig. 2d), indicating that vinculin was required for the subpopulation of integrins transmitting loads >7 pN (Fig. S7, S8). Thus, linkages to F-actin, but not integrin usage, could strongly influence the distribution of loads experienced by individual integrins.

We next examined the dynamics of force transmission at individual MTSs. We observed that, for a variety of cell types, the majority of load-bearing MTS_low_ and MTS_high_ molecules (>2 pN for MTS_low_; >7pN for MTS_high_) remained bound for up to tens of seconds at a close to constant force (Fig. 1d, Fig. 3b, Table S2). This observation is in apparent contradiction with previous formulations of the clutch model, which predicts continuous load-and-fail dynamics stemming from the progressive loading and failing of connections to F-actin (Fig. S22). Importantly, a minority of MTS FRET traces did exhibit anticorrelated changes in FRET donor and acceptor intensities indicative of dynamic changes in load (Table S3-S5, Fig. S9-S16). We have classified these as either step (close to instantaneous at our time resolution of ~1 second) or more gradual ramp transitions (Fig. 3a, 3c). Although the clutch model predicts ramp increases and step decreases in load, it does not predict the step increases or ramp decreases in load we observed (Fig. S9-S16). Although we cannot exclude the possibility that a subset of these events may be due to dye blinking, we rarely observe such events in our no-load control measurements (Table S6-S7). Measurements in U2OS osteosarcoma cells, which have been extensively used in studies of cell migration *(26)*, did not show step or ramp transitions (Fig. S17). Thus, dynamic load transitions, at least as assayed here, were dispensable for cell adhesion and migration.

**Fig. 3.**
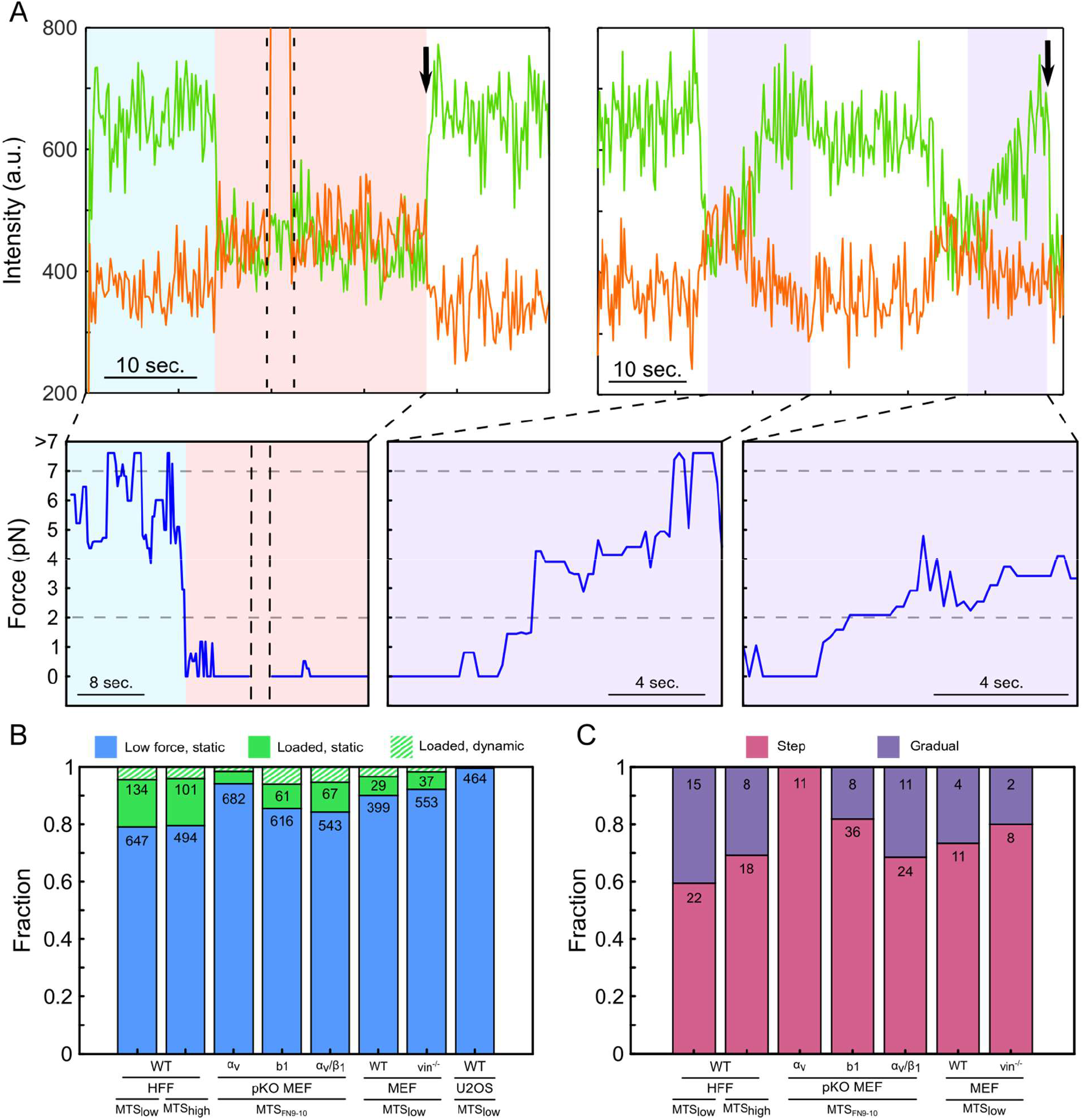
Dynamic transitions in load constitute a majority of sensor measurements. (A) Representative traces showing step (left) and ramp (right) load transitions (FRET donor: green; FRET acceptor: orange; load: blue) for HFFs adhering MTS_low_. Black arrows mark acceptor or donor bleaching; dashed black lines indicate direct excitation of the FRET acceptor. Horizontal gray dashed lines indicate upper and lower force measurement limits for MTSl ow. (B) Fraction of low force (defined as <2 or <7 pN) (blue), loaded (green), and dynamic (hashed; subset of loaded integrins) sensors for a variety of cell types adhering to different MTSs. (C) Fraction of sensors with step (magenta) and ramp (purple) transitions. U2OS cells had no observable dynamic events.

Our observations prompted us to explore alternative formulations of the clutch model (see Model Comparisons). Ultimately, we found that reversible crosslinks between actin filaments, for example mediated by α-actinin, allowed load to equilibrate across anchors, leading to long periods of close-to-constant loads (Fig. 4a-d). This updated model also yielded occasional step transitions that reflected the disconnection or reattachment of an individual anchor to an actin filament, and ramp transitions whose timescale reflected the equilibration of loads within the crosslinker network. In contrast, irreversible crosslinks between F-actin resulted in load-and-fail dynamics, analogous to previous clutch models (Fig. 4c). Such load-and-fail cycles are predicted to be costly in terms of energy dissipated in the repeated stretching of individual anchor linkages: in line with this understanding, models featuring reversible crosslinker dynamics predicted markedly lower energy dissipation for similar overall force levels (Fig. 4e).

**Fig. 4.**
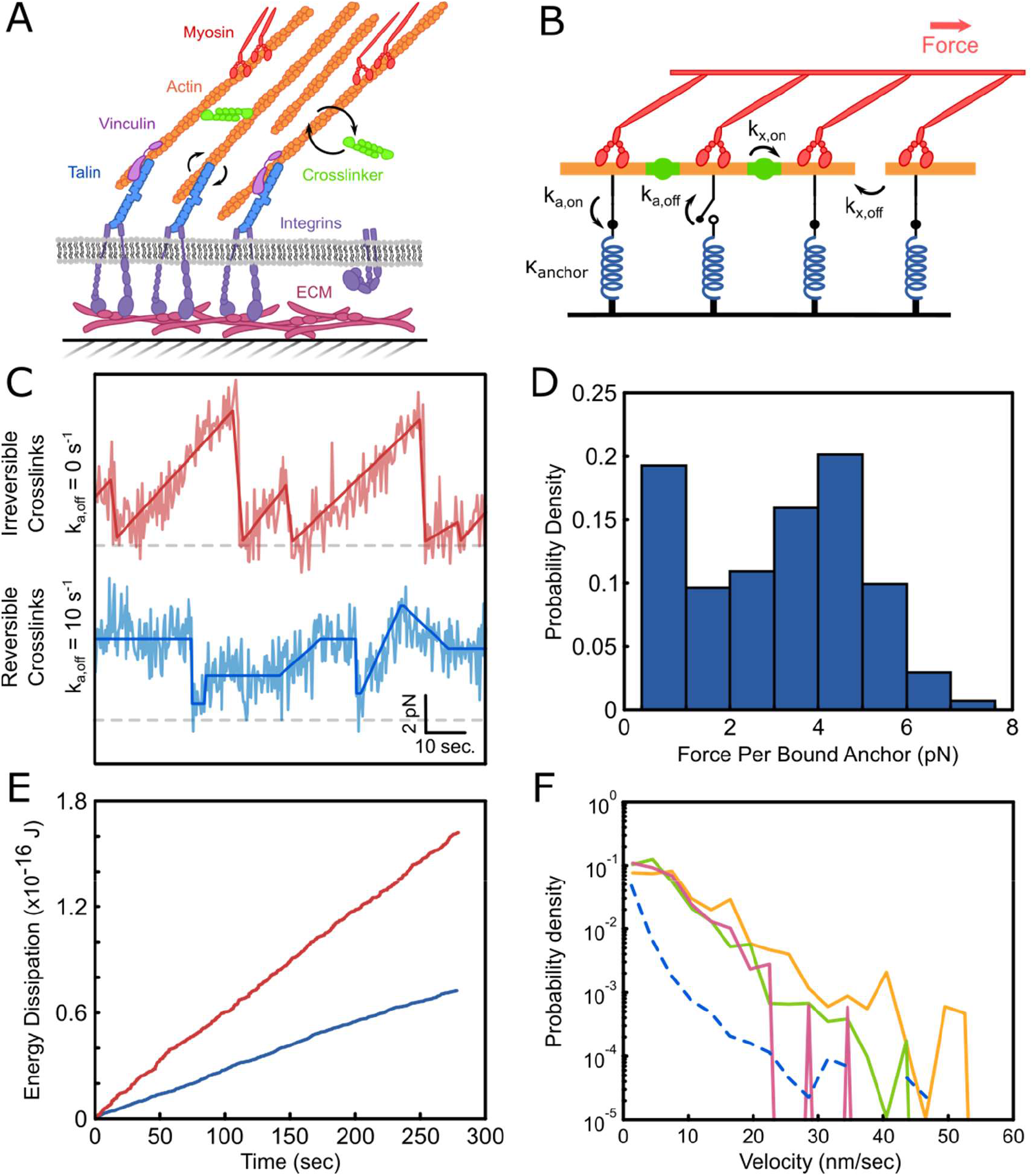
A modified model of cytoskeletal force transduction yields mechanical equilibrium at individual integrins. (A) Simplified cartoon of a focal adhesion: Nonmuscle myosin II pulls on reversibly crosslinked actin filaments, which are linked to integrins by vinculin and talin. (B) Cytoskeletal dynamics model: actin filaments bind to anchors (blue) and are linked by crosslinking proteins (green). (C) An example force trace of an anchor with irreversible (red) or reversible (blue) crosslinking. Reversible crosslinks allow for stable force plateaus as well as sporadic ramp and step events. (d) Force distribution for simulated anchors with reversible crosslinking (k_x,off_ = 10 s^−1^). (E) Calculated energy dissipation from simulations with irreversible (red) and reversible (blue) crosslinks. (F) Actin velocities measured for HFFs for tracks originating from segmented actin structures only (green), paxillin only (yellow), or overlapping paxillin and actin regions (pink). Dashed blue line: velocities predicted by simulations with reversible crosslinks.

The above model describes the loads borne by individual linkages to actin, rather than by the integrins themselves. Talin contains 3 actin binding sites, and can recruit up to 11 vinculin molecules *(27, 28)*. It is plausible that these connections to F-actin can act in parallel, potentially resulting in a broad range of loads transmitted by individual integrins. This possibility is supported by our observation of distinct subpopulations of integrins bearing ~6 pN and >10 pN within the FAs of HFFs (Fig. 1f). Single-pN loads are broadly consistent with a report that the average load experienced by talin was <6 pN (29). However, previous reports also describe >11 pN loads for a subset of talin molecules *(30)*, and peak loads of >50 pN for a subset of integrins *(21)*; these higher loads plausibly reflect additional, vinculin-mediated connections to F-actin (Fig. 2c, d). In total, these observations are consistent with the hypothesis that “digital” addition of discrete linkages to F-actin can result in a wide range of loads on individual integrins.

As an independent test of our model, we measured the velocity of individual F-actin filaments by treating living HFFs with 50 nM of SiR-actin, a fluorogenic small-molecule probe that binds to F-actin. The mean speed for F-actin within both adhesions and linear F-actin-rich structures (e.g. stress fibers) was ~5.5 nm/s, reasonably close to the mean velocity of 2.9 nm/s observed in simulations (Fig. 4f). This velocity is approximately one-third the magnitude of F-actin speeds measured in the lamellipodia of *Xenopus* XTC cells, a difference that may reflect a decrease in F-actin velocities near adhesions *(31)*. Simulations produced a power-law distribution of F-actin velocities that was qualitatively similar to experimental observation, providing independent evidence that the model could capture the essential aspects of force transduction.

Previously, we found that the majority of integrins exist in a minimally tensioned state *(14)*. Here we extend this result and report that a small fraction of ligand-engaged integrins support loads >11 pN. The large majority of integrins thus experience loads substantially less than their maximum capacity. This mechanical reserve may allow cells to withstand external stresses that would threaten tissue integrity. Conversely, the ability to exert large, localized forces via a few integrins may be essential for cell migration and mechanosensing, for example in fibrous ECM networks, where local effective stiffnesses can span several orders of magnitude *(32, 33)*. Integrin complexes thus represent an interesting example of how a highly asymmetric distribution of activity at the molecular level can yield flexible and robust functionality at the cell and tissue levels.

A core result of systems biology is that cellular subsystems, for example signal transduction pathways, are often organized into semi-autonomous functional modules, an outcome thought to enhance both robustness and evolvability *(34, 35)*. Though previously proposed *(6)*, whether a similar functional modularity might apply to complex structural assemblies such as FAs has been unclear. Our observations suggest that, despite a dense web of protein-protein interactions *(36)*, FAs maintain modularity at a functional level. In the model systems studied here, a common, albeit complex, force-transducing machinery resulted in per-integrin load distributions that were essentially identical regardless of integrin heterodimer usage (Fig. 2b). Integrin heterodimer usage in turn determined both ligand specificity and adhesion size, and hence cellular adhesion and traction output. The flexibility afforded by this modular organization is likely to have greatly facilitated the evolution of the remarkable functional diversity of integrin-based adhesion complexes in complex metazoans.

Our findings complement work demonstrating that some proteins are recruited to FAs as part of preassembled complexes *(37–39)*, suggestive of a hierarchical assembly process. Such preassembled protein complexes are however not necessarily synonymous with single, defined functions; indeed, in other systems evolutionary data demonstrate that biological function is often preserved even when the protein(s) fulfilling that function are not *(34)*. Structural and functional modularity may thus constitute distinct, and complementary, principles that govern the form and function of complex macromolecular assemblies.

Contrary to expectation, the majority of integrins experienced nearly constant loads within the resolution of our measurements *(7)*. Although several non-exclusive factors, for example domain unfolding in talin *(28)*, may contribute to this observation, a model that incorporates reversible crosslinks in the actin cytoskeleton is sufficient to account for our observations. This model is consistent with reports demonstrating that α-actinin crosslinking activity can change the mechanical properties of actin *(40)* and influence cell migration and traction force generation *(41)*. Force transmission through a network of dynamic crosslinkers also reduced energy consumption compared to a system that underwent repeated load-and-fail cycles (Fig. 4e). We therefore suggest that cellular mechanical homeostasis and efficient force transmission can potentially arise in a simple way from the core dynamical properties of the cytoskeleton.

## Acknowledgments

The authors thank K. Rothenberg of the Hoffman lab (Duke University) for the vinculin-null MEFs and M. Franklin from the Liphardt lab (Stanford University) for the U2OS cells. We thank the Khosla lab for their protein purification expertise and access to their equipment. We would also like to thank A. LaCroix from the Hoffman lab (Duke University) for useful discussions on FRET-force calibrations, and W. Weis and O. Chaudhuri for useful discussions and feedback. The data reported in this paper are further detailed in the supplementary materials.

## Author Contributions

S. J. T., A. C. C., and C. M. M. performed experiments and analyzed data, S. M. A. and L. S. P. created and ran clutch model simulations, all authors contributed to the writing and editing of the manuscript.

