## Supplementary Information for "Regulation and dynamics of force transmission at individual cell-matrix adhesion bonds"

**This PDF file includes:**

Materials and Methods

Supplementary Text

Figs. S1 to S26

Tables S1 to S9

### Materials and Methods

Sensor Construct Design

$MTS_{low}$ and $MTS_{SYN}$ were prepared as previously described (*14*, *15*). The high-force molecular tension sensor ($MTS_{high}$) was adapted from $MTS_{low}$ by replacing the $(GPGGA)_8$ module with another tension sensitive domain, termed $HP_{st}$, (LSDED FKAVF GMTRS AFANL PLWKQ QALMK EKGLF) derived from the the villin headpiece (*16*). The DNA encoding this construct was assembled by Epoch Life Sciences (Missouri City, TX) and was cloned into the pJ414 expression vector (DNA 2.0). We used Alexa 546 maleimide (ThermoFisher) as the FRET donor and an Alexa 647 maleimide dye (ThermoFisher) as the FRET acceptor. This modified MTS presents the identical RGD ligand derived from fibronectin as used in $MTS_{low}$. The entire $MTS_{high}$ sequence is presented below:

M G S E I G T G F P F D P H Y V E V L G E R M H Y V D V G P R D G T P V L F L H G N
P T S S Y V W R N I I P H V A P T H R S I A P D L I G M G K S D K P D L G Y F F D D H V
R F M D A F I E A L G L E E V V L V I H D W G S A L G F H W A K R N P E R V K G I A F
M E F I R P I P T W D E W P E F A R E T F Q A F R T T D V G R K L I I D Q N V F I E G T
L P M G V V R P L T E V E M D H Y R E P F L N P V D R E P L W R F P N E L P I A G E P
A N I V A L V E E Y M D W L H Q S P V P K L L F W G T P G V L I P P A E A A R L A K S
L P N A K A V D I G P G L N L L Q E D N P D L I G S E I A R W L S T L E I S G G A G E F
K C A G L S D E D F K A V F G M T R S A F A N L P L W K Q Q A L M K E K G L F G K C
A G S E N L Y F Q G T V Y A V T G R G D S P A S S A A H H H H H H

HaloTag domain
Dye labeling sites
Villin headpiece
TEV-cleavage site
RGD sequence from fibronectin

Expression and purification of MTS constructs

Sensors were expressed in BL21(DE3) competent *E. coli*. 500 mL cultures were grown overnight at 30 °C with 100 µg/mL ampicillin and induced with 1 mM IPTG at an OD of 0.6. The bacteria were then spun down at 6000xg for 30 min and resuspended in 10 mL lysis buffer (50 mM sodium phosphate, 300 mM NaCl, 10 mM imidazole, pH 8) with a protease inhibitor cocktail (11873580001 Roche) and 10 µM lysozyme. The resuspended cells were rocked for 30 minutes at 4°C, lysed with a tip sonicator, and spun at 14000xg for 30 minutes. The supernatant was incubated with 2mL of Ni-NTA HisPur Resin (Thermofisher) and rocked at 4 °C for 2 hours. The solution was then packed into a gravity column, washed with 3 times with 5 mL of Wash Buffer (50 mM sodium phosphate, 300 mM NaCl, 20 mM imidazole, pH 7.4), and incubated with 4 mL of Elution Buffer (50 mM sodium phosphate, 300 mM sodium chloride, 250 mM imidazole, pH 7.4) for 5 minutes. The eluate was collected and dialyzed overnight into Storage Buffer (1x PBS, 1 mM EDTA, and 2 mM β-mercaptoethanol), flash frozen, and stored at -80 °C. Fractions were characterized by SDS-Page and the concentration was determined by UV-Vis spectroscopy.

Labeling of MTS constructs

Labeling of MTSs was done through dual cysteine labeling and subsequent purification to separate the population of sensors with a single donor and acceptor dye. The cysteines were first reduced with 2 mM TCEP for 30 minutes at room temperature and buffer exchanged into Labeling Buffer (50 mM phosphate buffer, 150 mM NaCl, 1mM EDTA, pH 7.4) using 3 7K Zeba desalting columns (89883, Thermofisher) in series. Alexa 546 and Alexa 647 maleimide dyes (Invitrogen) were added at a protein:donor:acceptor ratio of 1:1.5:2 for 1 hour at room temperature and overnight at 4°C. To help remove free dye and exchange the protein into FPLC buffer A (50 mM Tris buffer, pH 8, 5 mM β-mercaptoethanol), the solution was passed through 2 PD Minitrap desalting columns (45001529, GE Healthcare) in series. To separate out the sensors with a single donor and single acceptor, we used an AKTApure FPLC (GE Healthcare) with a MonoQ PC 1.6/5 (GE Healthcare) ion exchange column and a 10 mM/mL linear salt gradient with Buffer B (50 mM Tris, pH 8, 5 mM β-mercaptoethanol, 2 M NaCl). Fractions were characterized using SDS-Page, UV-Vis spectroscopy, and single-molecule imaging. The desired fractions were concentrated and exchanged into PBS using 3K centrifugal filters (Amicon), and stored at -80°C.

Preparation of functionalized HALO ligand coverslips

Coverslips were prepared as previously described (*14*). Briefly, 24x50 mm No. 1 coverslips (Fisherbrand) were sonicated in a bath sonicator (Kendall) for 20 min with isopropanol, MilliQ water, and 5 M KOH, with MilliQ water rinses between each step. The coverslips were then sonicated for 5 minutes in methanol and transferred to a solution of 2 mL N-(2-aminoethyl)-3-aminopropyltrimethoxysilane (97%) (A0700, UCT Specialties), 10 mL glacial acetic acid, and 200 mL methanol. The coverslips were incubated in the silane mixture for 10 minutes, sonicated for 1 minute, and then incubated for another 10 minutes. They were then rinsed with MilliQ water and dried with nitrogen. To passivate the coverslips, 100 mg of maleimide PEG Succinimidyl Carboxymethyl Ester, MW 5000 (A5003-1, JenKem Technology) was dissolved in 1 mL of 100 mM phosphate buffer (pH 7.0). Two coverslips were sandwich with 100 μL of the PEG solution in between for 1 hour at room temperature and protected from light. The coverslips were then washed with MilliQ water and dried before being incubated overnight with 100 μL of 3 mM Halo ligand thiol (P6761 Promega) in 100 mM phosphate buffer (pH 7.0). Afterwards, the coverslips were washed with MilliQ water, dried, and stored in vacuum-sealed bags at -20°C.

Flow Chamber Preparation and Imaging

Flow chambers were attached to PEGylated coverslips as previously described (*15*). For ensemble experiments, chambers were prepared with 100 nM of double-labelled sensor and incubated at room temperature for 30 min. For the single-molecule assay, 100 nM of unlabeled sensor with 100 pM of labeled sensor was mixed in PBS and added to the flow cells for 30 min. The chambers were then washed with 200 μL PBS and Pluronic F- 127 (0.2% w/v) for ~1 minute to prevent non-specific cell attachment. The chambers were washed again with PBS to remove excess Pluronic. Cells were then added and incubated for at least 1 hour at 37 °C in DMEM high glucose medium. FRET measurements were made within 3 hours of plating the cells and acquired with an objective heater (Bioptechs) set to 37 °C. Images were prepared in Fiji (*43*) and analyzed using custom Matlab scripts.

TIRF FRET imaging

Single-molecule and ensemble FRET fluorescence measurements were performed with objective-type total internal reflection fluorescence (TIRF) microscopy on an inverted microscope (Nikon TiE) with an Apo TIRF 100x oil objective lens, NA 1.49 (Nikon) as described previously (*14*) and controlled using Micromanager (*44*). Samples were excited with 473 nm Obis laser (Coherent), 532 nm (Crystalaser), or 635 nm (Blue Sky Research) lasers. For single-molecule data, emission for the FRET donor and emission channels were separated as previously described and recorded on an EMCCD camera (Andor iXon) (*15*). For collection of the GFP signal, we used an additional set of emission filters mounted on a motorized flip mount (Thor labs) placed the donor fluorescence emission path. Filters used included a 593 nm/40 nm filter (Semrock) for the collection of donor emission, a 675/30 nm filter for the collection of acceptor emission, and a 514 nm/30 nm filter (Semrock) for GFP emission collection. For ensemble FRET maps taken for whole cells, emitted light passed through a quad-edge laser-flat dichroic with center/bandwidths of 405 nm/60 nm, 488 nm/100 nm, 532 nm/100 nm, and S5 635 nm/100 nm from Semrock (Di01-R405/488/532/635-25x36) and corresponding quad-pass filter with center/bandwidths of 446 nm/37 nm, 510 nm/20 nm, 581 nm/70 nm, 703 nm/88 nm band-pass filter (FF01- 446/510/581/703-25). GFP, donor, and acceptor images were taken through separate additional cubes stacked into the light path (GFP: 470 nm/40 nm, 495 nm LP, 525 nm/50 nm; donor: 550 nm LP; acceptor: 679 nm/41 nm, 700 nm/75 nm) and recorded on a Hamamatsu Orca Flash 4.0 camera.

Cell Culture

Human foreskin fibroblast (HFF) cells CCD-1070Sk (ATCC CRL-2091) were cultured in DMEM high glucose medium (Gibco, Cat #21063-029) in the absence of phenol red and supplemented with 10% fetal bovine serum (FBS, Axenia Biologix), sodium pyruvate (1 mM, Gibco), MEM non-essential amino acids (1x, Gibco), and penicillin/streptomycin (100 U/mL and 100 μg/mL, Gibco), herein referred to as normal culture media. The cells were grown at 37 °C with 5% $CO_2$. Fibroblasts with stably expressing GFP-paxillin at the C-terminus were prepared as previously described (*15*).

Integrin pan-knockout (pKO) mouse kidney fibroblasts rescued with either $\alpha_v$, $\beta_1$, or both $\alpha_v$ and $\beta_1$ integrin subunits, were a generous gift from Reinhard Fässler (MPI Martinsried) (*22*). Cells were cultured on fibronectin-coated plastic (5 μg/mL, Corning, diluted in PBS and incubated at 37 °C for 1 hr) in normal culture media described above. pKO-β1 cells in particular were sensitive to the quality of fibronectin coating, thus a minimum of 1 mL and 4 mL of the diluted 5 μg/mL fibronectin solution were used per well for a 6-well and 10 cm dish, respectively. Cells were grown at 37 °C with 5% $CO_2$.

Wild-type and $vin^{-/-}$ mouse embryonic fibroblasts were a generous gift from Brent Hoffman (Duke University) (*45*). Cells were cultured on tissue culture plastic in normal culture media at 37 °C and 5% $CO_2$.

U2OS cells were a generous gift from Jan Liphardt (Stanford University). Cells were cultured on tissue culture plastic in normal culture media at 37 °C and 5% $CO_2$.

Transfection

pKO-integrin MEF cells and wild-type and vinculin KO MEF cells were transfected using a similar protocol to the previously described GFP-paxillin human fibroblasts (*15*). Cells were trypsinized, pelleted, resuspended in media lacking FBS and penicillin/streptomycin, and counted. 2x10$^6$ pKO-integrin cells and 5x10$^5$ wild type and vinculin KO cells were re-pelleted at 800 rpm for 10 min. 82 µL of P4 Nucleofector solution was added to 18 µL of P4 supplement in a 1.5 mL Eppendorf tube and used to resuspend the cell pellet. DNA for C-terminus GFP-paxillin (Addgene #15233) cloned into the DNA 2.0 PiggyBac vector (~4 µg) was added to the cells and gently flicked before transferring to a Lonza nucleofection cuvette. Cuvettes were placed in a Lonza 4D-Nucleofector system and program C2167 (for mouse embryonic fibroblasts) was used. 500 µL of warm media was added to the cuvette and cells were transferred to a 6-well plate with media equilibrated at 37 °C with 5% CO2 using a pipet bulb without pipetting up and down. Cells were selected with 1-2.0 µg/mL puromycin 24 h after transfection, for 4-6 days.

Calculating single-molecule FRET efficiency

Single-molecule data were acquired and analyzed as described previously (*14*). Briefly, data were acquired with excitation with a 532 nm laser at 5 frames per second for 300 or 600 frames and with direct acceptor excitation at 635 nm for approximately 10 frames at roughly frame 100. The direct excitation helped to distinguish between low FRET sensors and sensors without an acceptor dye.

Traces were analyzed using a custom Matlab code and donor and acceptor channels were aligned using a SHREC (Single molecule High Resolution Colocalization) map generated by scanning across a field of beads (*46*). The positions of individual sensors were then detected using a spot-finding algorithm (Tristan Ursell, Stanford University) and were determined to be co-localized if within 2 pixels. Intensities were calculated based on an average of 7x7 pixels centered around the detected spot and corrected for spectral bleedthrough.

Intensities for each dye were averaged over manually identified FRETing, non-FRETing, and bleached regions. When the acceptor bleached before the donor, we used the following expression to calculate FRET efficiency:

$$E = \frac{(I_a - I_{a,back})}{(I_a - I_{a,back}) + \gamma(I_d - I_{d,back})}$$

$$\gamma = \frac{I_a - I_{a,back}}{I_{d,0} - I_d}$$

or equivalently,

$$E = \frac{I_{d,0} - I_d}{I_{d,0} - I_{d,back}}$$

Where:

$I_a$ = acceptor intensity during FRET

$I_{a,back}$ = acceptor background intensity

$I_d$ = donor intensity during FRET

$I_{d,0}$ = donor intensity after acceptor photobleaching

$I_{d,back}$ = donor background intensity

$\gamma$ = correction factor accounting for relative dye quantum yields and instrument detection efficiencies

When the donor fluorophore bleached first, the FRET efficiency was calculated as:

$$E = \frac{(I_a - I_{a,back})}{(I_a - I_{a,back}) + \gamma_0(I_d - I_{d,back})}$$

Values for $\gamma_0$ were 0.40 for $MTS_{low}$, 0.52 for $MTS_{FN}$, and 0.52 for $MTS_{high}$. Events were double-checked by generating a series of $z$-projections for the donor and acceptor molecule during FRETing, non-FRETing, and bleached states. The autoGaussianSurf Matlab function (Patrick Mineault) was used to fit a 2D Gaussian to the 7x7 pixel area to determine if the spot represented a single emitting fluorophore. Low FRET events were verified as having a functional acceptor by directly excitation with a 635 nm laser.

Obtaining theoretical FRET-force calibration curve

The FRET vs. force response of the $(GPGGA)_8$ linker used here was previously reported by Grashoff et al., and an updated calibration was recently reported by LaCroix et al. (*42*, *47*). We used the updated Matlab calibrations from LaCroix et al. to generate improved FRET vs. force calibration curves. Using 43 amino acids (for the 8 repeats of GPGGA plus the two cysteines and a single lysine), a fluorophore radius of 0.5 nm, a Forster radius of 6.95 nm, and persistence lengths from 0.87 to 0.98 nm, we constructed 3 FRET-force calibration curves to account for the slightly different resting FRET efficiencies determined experimentally (Fig. S18).

Ensemble FRET analysis

Ensemble measurements were performed as previously described (*14*). In summary, images of GFP-paxillin marked cells were acquired using a Hamamatsu Orca Flash4.0 camera, and were subsequently corrected for illumination spatial inhomogeneities, background subtracted, and intensity normalized. The GFP image was then boxcar averaged ("moving average v3.1" from MATLAB Central File Exchange) at 10 different rotations of the original image at 20° intervals, thresholded, and segmented using a watershed algorithm. The segmented image was then corrected to combine adjacent islands representing a single adhesion and filtered to exclude islands below a lower limit (0.5 $\mu m^2$). The segmented GFP image was then used to mask the corresponding FRET signal.

FRET images were converted to FRET index values by dividing the acceptor intensity over the sum of the donor and the acceptor signal. Then, the FRET images were converted to FRET efficiency after correcting to dye labeling efficiency, bleedthrough, the measured no-load FRET efficiency, and the FRET-index measured outside the cell (*14*). The total force of a cell was calculated by summing the force contributions (defined as the average pixel value times number of pixels) for pixels within adhesions. For $MTS_{high}$ measurements, calculated forces corresponding to <7 pN were set to 0 pN.

Dynamics analysis

Traces with potential dynamic behavior, identified by having distinct anti-correlated signals, were marked and analyzed individually. Step events were classified by having large, anti-correlated changes in the donor and acceptor signal within the period of 1-3 frames (0.2-0.6 seconds) and were manually annotated. Ramp events were classified by having more gradual anti-correlated changes and were manually marked. Ramp traces were then converted to the force domain and only changes in load between 2 and 7 pN were fit. Dynamic events were only accepted if the acceptor was confirmed to be active, and could not be accounted for by either donor or acceptor photobleaching.

Actin tracking imaging

Halo-PEG coverslips were incubated with 100 nM unlabelled RGD sensor at RT for 30 min, and cells were seeded and allowed to spread for at least ~1 hr. After a 9 minute incubation with 50 nM SiR-Actin (Cytoskeleton Inc., #CY-SC001), the sample was incubated with Prolong Live Antifade Reagent (Invitrogen P36975) for 1 hr. The low dilution of SiR-Actin allowed for individual molecules to be tracked, and the addition of Prolong reduced photobleaching. For each cell, a 100 ms exposure of the GFP-paxillin channel (for masking) was first acquired, followed by a 60 frame sequence in the red channel (SiR-Actin) with 300 ms exposures taken every 2s.

For fixed cell data, cells were allowed to spread on functionalized coverslips and then fixed in 4% paraformaldehyde for 15 minutes at room temperature. After rinsing, the cells were treated with SiR-Actin and Prolong reagent as above.

Actin tracking analysis

Speckles and tracks were identified using QFSM software made available by the Danuser lab (*48*) (Fig. S19). The localizations, which are given with pixel precision, are fit to subpixel positions by Gaussian fitting. The $x$ and $y$ positions of the points in each track were independently fit to linear fits of $x$ vs. time and $y$ vs. time, and the slopes of these fits gave the velocity of each track. Errors for each velocity were propagated from the standard errors of both slopes.

Velocity histograms of actin within cell regions include all tracks whose first point originates within a given mask. The total cell area was masked using a mean threshold of the GFP-paxillin image, where cytoplasmic paxillin signal was used to segment the cell. After dilating and eroding to fill small holes, the edge of this mask was smoothed by fitting to a cubic spline curve. Adhesions were masked by an Otsu threshold of the same GFP-paxillin image after background subtraction to remove the diffuse cytoplasmic signal. Actin stress fibers were masked as the brightest 2% of pixels of a time-series projection of the actin tracks. Finally, the cytosol was taken as regions within the cell mask which are excluded from both the actin mask and the paxillin mask.

We suspected the measured velocities in live cells could reflect a convolution of some noise function and the true velocities. To derive an empirical estimate of the noise function, we acquired actin tracks in paraformaldehyde fixed cells exactly as described above and used the velocities of these tracks to determine the measurement noise. When separated by the number of points in a track (called track length hereafter), it was apparent that the fixed cell velocity distributions narrow and shift toward zero with increasing track length, indicating that the measurement noise is a function of the number of points in a track (Fig. S20). Therefore, we deconvolved each velocity distribution for a given track length separately. In short, a guess for the true velocity pdf, $g(v)$, for a given track length (originally taken to be the measured distribution, $f(v)$) is convolved with the measured fixed-cell pdf, $n(v)$, for the same track length. The true velocity pdf $g(v)$ is then optimized by non-linear least squares minimization of the difference $g(v)*n(v) - f(v)$ where * is the convolution operator. This optimization is done in Matlab using the trust-region-reflective algorithm which accepts upper and lower bounds on the solution, keeping probabilities between zero and one. The total pdf is then reconstructed by summing each deconvolved pdf for a given length, weighted by the number of tracks with that length. All pdfs are estimated by the histograms of the velocity distributions with a bin width of

3nm/s.

Actin tracks were simulated to verify under what conditions deconvolution provides an accurate estimate of true velocity. For a given actin population (e.g. live or fixed cell), a velocity distribution was chosen to represent the 'true' actin velocities. A track length distribution was also set to approximate that of the measured data, which is long-tailed and well-fit by a Pareto distribution with $x_m = 4$ (for tracks of length >= 4) and $\alpha = 1.97$ (in number of frames). From these two distributions, individual tracks were generated, each with a given velocity and track length. To recapitulate the actual measurements, where track localizations are defined by cartesian coordinates, a random angle was generated between 0 and $2\pi$ and the velocity split into $x$ and $y$ components according to this angle. The "true" track is then modelled as a constant velocity track with localizations linearly spaced from the origin to the end point determined by the velocity and track length. Noise was added to these localizations, which followed a logistic distribution with $\mu = 0$ and $s = 0.75$ (in pixels). Velocities were then calculated from these localizations using the same analysis pipeline as the original data.

For the fixed cell simulation, true velocities were always zero. With the addition of noise, the velocity distribution for each track length qualitatively matched the measured distribution in fixed cells, indicating the noise model is an appropriate choice to represent our data (Fig. S21). For the live cell distribution, the true velocity was guessed to follow a long-tailed gamma distribution with $k = 1.5$ and $\theta = 1.8$ (in nm/s), which, after noising, approximated the measured actin velocities in living cells over focal adhesions.

After simulating these two distributions, they are put through the same deconvolution process described above. This procedure revealed that only at longer track lengths (≥13 localizations) does the deconvolution yield accurate results (Fig. S22), indicating that for shorter tracks, the measured velocity is not well-described by a simple convolution of the fixed cell velocity distribution and the true velocity distributions. Therefore, only the tracks with at least 13 localizations are included in the velocity histograms.

### Supplementary Text

Lifetime Analysis

We assume any given sensor under load can undergo three competing terminations: either acceptor bleaching ($k_{Abl}$), donor bleaching ($k_{Dbl}$), or integrin unbinding ($k_I$). To provide a simple estimate of bound-state lifetimes for the entire pool of integrins, we assume first-order kinetics and that the lifetimes are insensitive to load and integrin type. This allows us to estimate the fraction of events that terminate due to integrin unbinding as the ratio of $k_I$ to the sum of all possible rates:

$F_{RI}: Fraction\ of\ events\ that\ end\ due\ to\ integrin\ unbinding$
$k_I: Rate\ of\ integrin\ unbinding$
$k_A: Rate\ of\ acceptor\ bleach$
$k_D: Rate\ of\ donor\ bleach$

$$F_{RI} = \frac{k_I}{k_D + k_A + k_I} \rightarrow k_I = \frac{F_{RI}(k_D + k_A)}{1 - F_{RI}}$$

We measured the average bleaching time for the donor ( $\tau_{Dbl}$) and acceptor ( $\tau_{Abl}$) for each of our MTSs under no-load conditions. (Since any given sensor is initially at a high FRET efficiency, we calculated the average donor bleaching time by measuring the donor lifetime after the acceptor had already bleached, for molecules in which acceptor bleaching happened first.)

| | $\tau_{Abl}$ (s) | $k_{Abl}$ ($s^{-1}$) | $\tau_{Dbl}$ (s) | $k_{Dbl}$ ($s^{-1}$) |
|---|---|---|---|---|
| $MTS_{low}$ – no-load | 12 | 0.083 | 38 | 0.026 |
| $MTS_{low}$ – no-load* | 11 | 0.089 | 23 | 0.043 |
| $MTS_{high}$ – no-load | 15 | 0.067 | 43 | 0.023 |
| $MTS_{FN9-10}$ – no-load | 19 | 0.054 | 37 | 0.027 |

*Denotes no-load measurements taken for WT and $vin^{-/-}$ MEF data sets with a new green laser.

From our single molecule measurements, we observed a number of events exhibiting a step decrease in force, indicative of integrin unbinding. To calculate the fraction of events terminating in unbinding, we take the ratio of observed step unloading events divided by the number of molecules bearing appreciable load (>3pN for $MTS_{low}$ and $MTS_{FN9-10}$ or >7pN for $MTS_{high}$). For human fibroblasts adhering to $MTS_{low}$, we observed 24 step unbinding events and 134 sensors bearing > 3pN. Using that as the fraction of events due to unbinding, we can calculate $k_I$ for HFFs adhering to $MTS_{low}$:

$$F_{RI} = 24/134 = 0.179$$

$$k_I = \frac{F_{RI}(k_D + k_A)}{1 - F_{RI}} = \frac{0.179(0.083\ s^{-1} + 0.026\ s^{-1})}{1 - 0.179} \rightarrow\ 1/k_I = 42\ s$$

Repeating a similar analysis for different cell and sensor combinations, we can calculate the average integrin lifetime for each condition (Table S2).

Model Comparisons

Prior to these experiments, the single-clutch dynamics predicted by the base motor clutch model as described originally by Chan and Odde, and later by Bangasser et al. and Mekhdjian et al., had not been tested experimentally (*2*, *9*, *12*). The original model predicts forces on single clutches rising approximately linearly (Fig. S23a), even in a "stalled" state (i.e. mean actin retrograde flow speeds near 0 nm/s), contradicting our experimental observation that indicate integrins exist in force equilibriums persisting for 10-30 seconds (Fig. 3A). In addition, the model predicts force distributions that decrease monotonically (Fig. S23b), though our results demonstrate distinct subpopulations of force. To account for these discrepancies, we explored modifications to the motor clutch model that could reproduce the characteristic our single molecule measurements.

Vinculin Reinforcement and Catch Bond

In attempt to explain these features of the load distribution and dynamics, we considered alternative models that incorporate either force-dependent clutch recruitment (*2*) or clutch unbinding following a catch bond model (*12*). To replicate the recruitment of additional, reinforcing linkages to actin, we constructed a model in which any clutch whose force rises above a force threshold representative of talin unfolding ($F_{talin}$) becomes less likely to unbind, modeled as an increase in bond rupture force ($F_b$) and a decrease in basal clutch unbinding rate ($k_{off}$). This allowed "force-activated" clutches to reach significantly higher force peaks, leading to high force events as observed experimentally (Fig. S24a, b). In order to develop force histograms with a peak, the clutches were modeled as a catch bond instead of a slip bond, leading to a force peak at 2-3 pN. Combination of reinforcement with a catch bond revealed a second peak at 6-7 pN. Though this accurately captured the force distributions observed, the dynamics at the single-clutch level were identical to the base model (Fig. S24a) and we did not observe any force plateaus.

<u>Viscous relaxation and rapid re-binding</u>

Next, we challenged the assumption of instantaneous clutch relaxation included in our previous models. If the clutches do not completely relax due to viscous resistance to elastic recoil, then the clutches could rebind at appreciable load and resume the loading phase. Given the noise in the experimental measurements, such fluctuations might be small enough to be consistent with our data. We modeled time-dependent viscous relaxation of the clutches as an exponential (Eq 1), and we assumed the binding rate of the clutches was dependent on its distance from the actin filament (Eq 2). However, the clutch traces still exhibited extremely dynamic load and fail behavior (Fig. S25) and did not reproduce the plateaus of forces observed.

$$x_{cl,t} = x_{cl,t-1}\left(e^{-\frac{K_c}{N}}\right)\Delta t \quad Eq. (1)$$

$$k_{on} = k_{on,basal}\left(e^{C\left(x_{cl,unbind} - x_{cl,t}\right)}\right) \quad Eq. (2)$$

<u>Multiple connections model</u>

The next model we explored aimed to better capture the experimental dynamics by including a more structurally-detailed approach to modeling reinforcement. When vinculin reinforces talin, it creates up to 11 additional connections between F-actin and integrin (*28*). To represent this effect, we developed a multiple connections model in which each clutch was composed of an integrin spring and a varying number of talin/vinculin springs connecting the integrin to actin (Fig. S26a). When the number of connections was low (2-4), a single talin/vinculin spring unbinding led to a large change in the force on the integrin (Fig. S26b). When many (6-12) connections between the integrin and actin were used, the force on the integrin spring remained relatively constant (Fig. S26c). Because the duration of the plateaus did not match what we observed experimentally, this model did not capture the force plateaus across conditions.

<u>Dynamic F-Actin Network</u>

The base model assumed F-actin to act as a single filament pulled by a cluster of myosin motors. In the dynamic F-actin model, the F-actin was treated as a network, connected by noncompliant dynamic crosslinkers (Fig. 4). These crosslinkers can bind and unbind at rates $k_{x,on}$ and $k_{x,off}$. Each clutch is bound to an individual filament, and the number of motors per filament is dictated by simulation parameters (Table S9). The stiffness of the substrate as defined in the base model is taken to be very high to match the glass surfaces used experimentally. Because the binding rate constant of the crosslinkers is 1-2 orders of magnitude higher than the unbinding rate constant, the clutches exist in clusters bound to the same filament. The motors pull the filaments, and the forces on individual clutches build until the simulation reaches a "stalled" state, meaning the F-actin retrograde flow rate is close to 0 and the motor stall force and clutch force are nearly equal. When a crosslinker unbinds the force on each filament is no longer balanced. Because the forces on the clutches were not necessarily evenly distributed, one of the new filaments can have a total clutch force greater than the stall force of the motors. This will cause the F-actin to briefly move in the opposite direction (anterograde flow) until a force balance is restored. The other filament will continue to move in the typical direction (retrograde flow) to restore force balance. In the base model, there is no mechanism for the development of anterograde flow, so even if the retrograde flow rate is very small, the force on clutches will continue to build. Without anterograde flow, the clutch cannot relax and the force cannot fluctuate at near-mechanical equilibrium.

Throughout the simulations, dynamic clusters of clutches continuously stretch and relax, oscillating at force plateaus for periods of 10-60 seconds. Ramp and step transitions are observed throughout these simulations, in a manner consistent with our experimental observation. A step transition occurs when a binding clutch quickly builds force, or when an unbinding clutch instantaneously returns to 0 force. When a clutch or crosslinker binds or unbinds, the neighboring clutches adjust to achieve a balanced force, resulting in a more gradual ramp transition for the attached integrins. In addition, the tendency of clutches to reach a force and plateau leads to peaked force distributions that can be altered by varying simulation parameters.

Though a variety of models were explored, the dynamic actin network best captured the behavior observed experimentally. In contrast to the other models, the force plateaus persisted the longest with minimal fluctuations, and the single-clutch dynamics were consistent across simulation parameters (i.e. relatively insensitive to small parameter changes). Although further testing of the dynamic F-actin network model is still needed, and alternate models have not been definitively ruled out, the dynamic F-actin model presented in the paper best captures the experimentally observed behavior of the individual sensor force distribution and force dynamics.

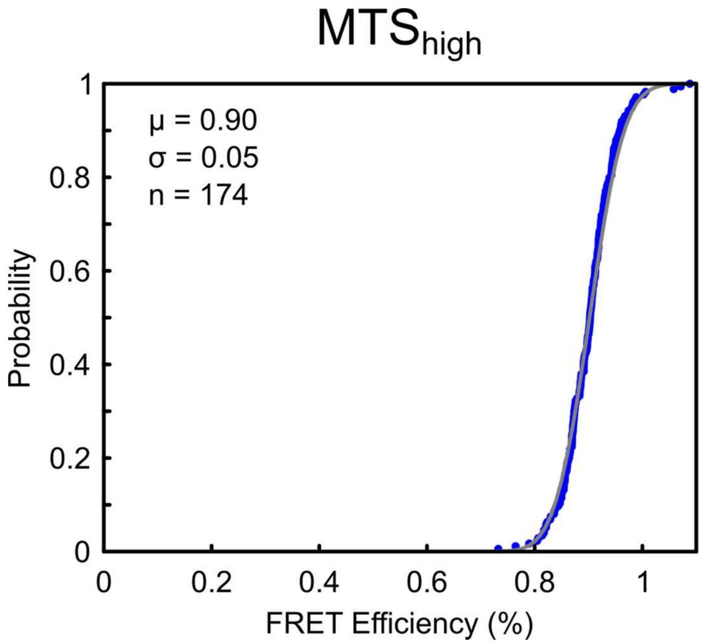


**Fig. S1. No-load FRET cumulative distribution for MTS$_{high}$.**

No-load single-molecule FRET measurements for MTS$_{high}$ with 1 hr incubation at 37 °C in normal cell culture media. Grey line indicates Gaussian fit with mean (μ) and standard deviation (σ).

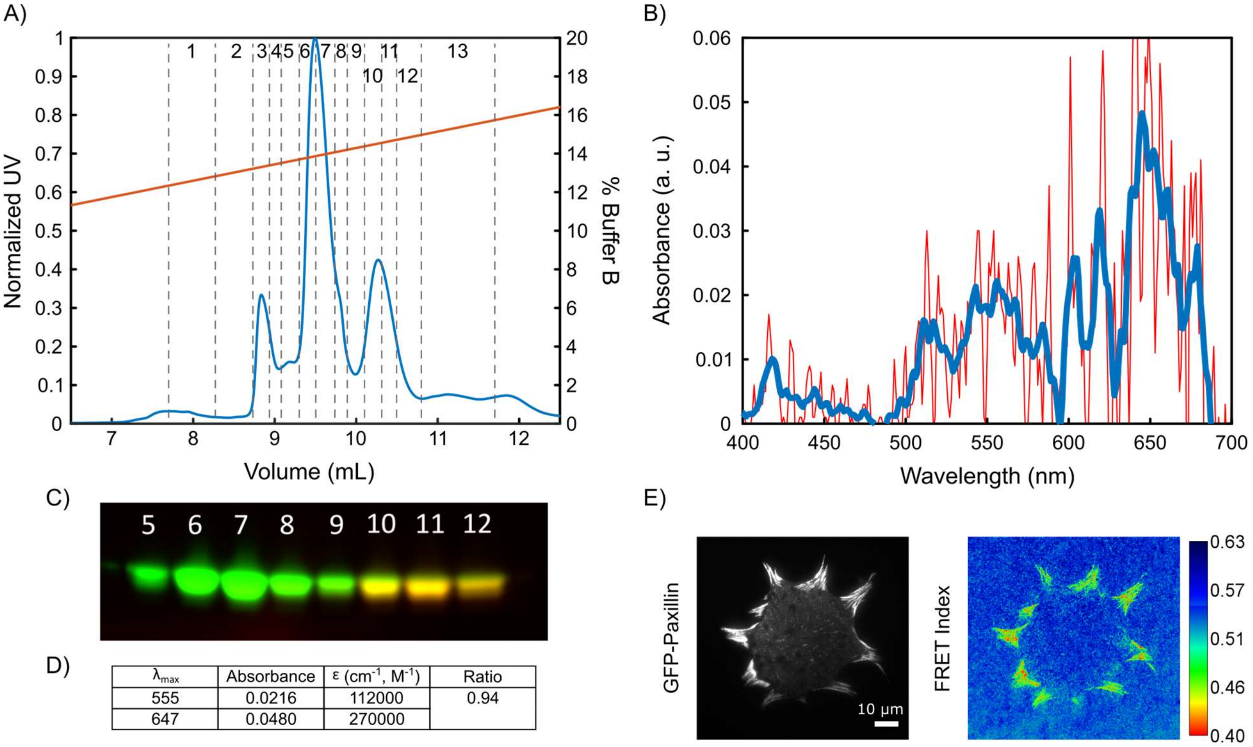


| $\lambda_{max}$ | Absorbance | ε ($cm^{-1}$, $M^{-1}$) | Ratio |
|---|---|---|---|
| 555 | 0.0216 | 112000 | 0.94 |
| 647 | 0.0480 | 270000 | |

**Fig. S2. Biochemical characterization of MTS$_{high}$.**

(A) FPLC trace with anion exchange column (MonoQ PC 1.6/5) of MTS$_{high}$ and accompanying UV (blue) and % Buffer B (orange) traces. Fractions were collected manually, as numbered and indicated by the vertical lines. Fractions 10 and 11 were combined for further experiments. (B) Absorbance trace from 400 to 700 nm for collected fractions with raw data (red) and moving average (blue). (C) SDS-PAGE gel of FPLC fractions fluorescently imaged with a Typhoon 9410 gel scanner. (D) Final quantification of labelling efficiency of collected FPLC fractions, measured with a 1mm Hellma Tray Cell on an Eppendorf BioPhotometer Plus spectrometer. (E) Ensemble image of GFP-Paxillin HFFs adhered to MTS$_{high}$.

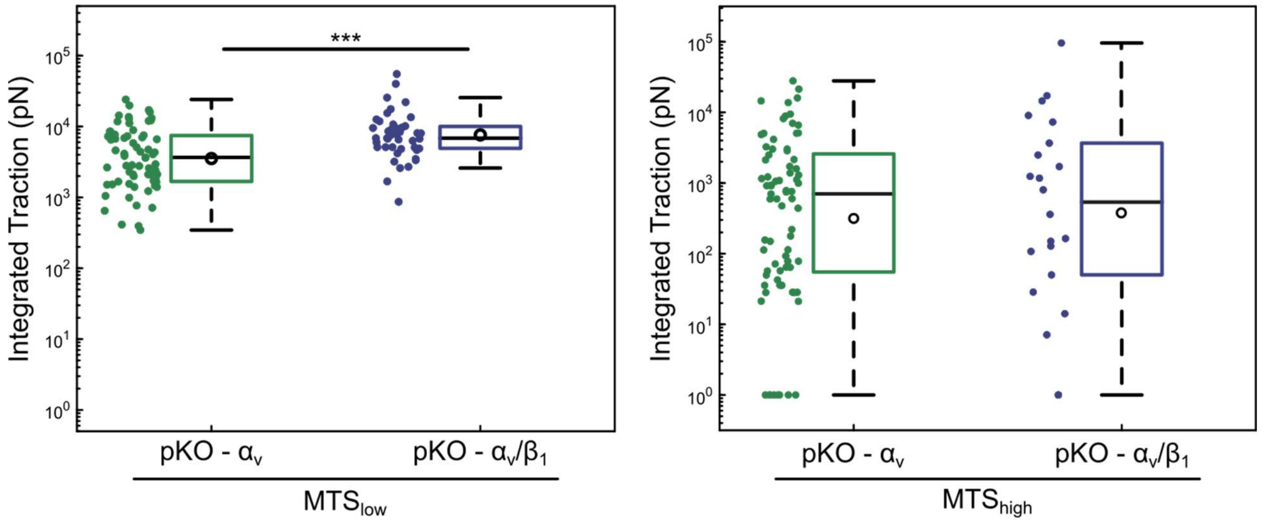


**Fig. S3. Ensemble quantification of pKO-$\alpha_v$ and pKO-$\alpha_v/\beta_1$ MEF cells adhering to MTS$_{low}$ and MTS$_{high}$.**

When adhering to MTS$_{low}$, pKO-$\alpha_v/\beta_1$ cells exert more integrated traction compared to pKO-$\alpha_v$ cells (pKO-$\alpha_v$:67 cells, Mean: 35.3 nN; pKO-$\alpha_v/\beta_1$: 43 cells, Mean: 72.5 nN). (***$p = 3x10^{-4}$) When adhering to MTS$_{high}$, pKO-$\alpha_v/\beta_1$ and pKO-$\alpha_v$ cells produce comparable traction overall (pKO-$\alpha_v$:77 cells, Mean: 2.6 nN; pKO-$\alpha_v/\beta_1$: 22 cells, Mean: 7 nN).

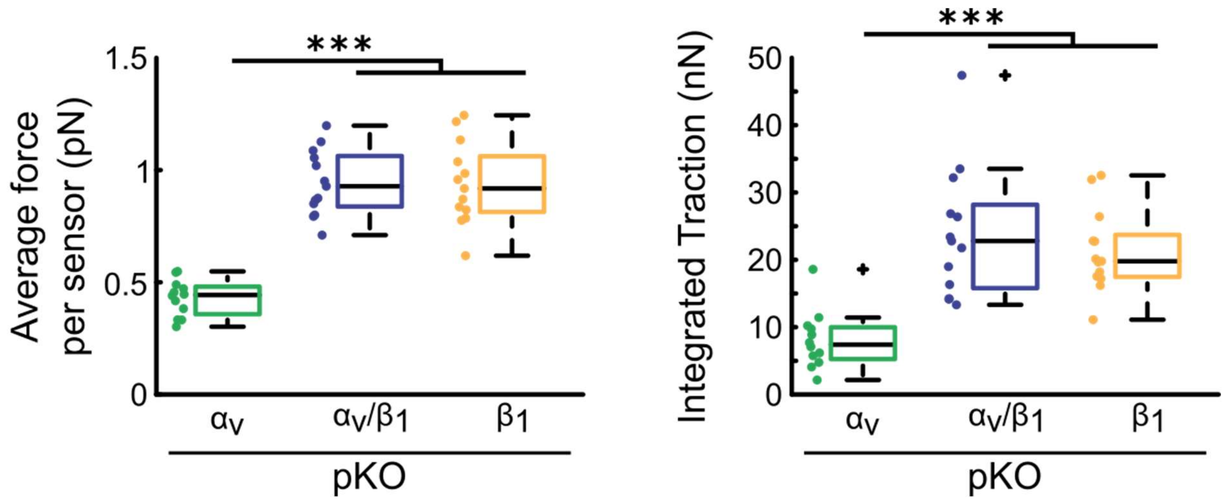


**Fig. S4. Ensemble quantification of pKO-$\alpha_v$, pKO-$\alpha_v/\beta_1$, pKO-$\beta_1$ MEF cells adhering to $MTS_{FN9-10}$.**

pKO-$\alpha_v/\beta_1$ and pKO-$\beta_1$ cells exert a higher average load per sensor and higher integrated traction as compared to pKO-$\alpha_v$ cells (pKO-$\alpha_v$:13 cells; pKO-$\alpha_v/\beta_1$: 12 cells, pKO-$\alpha_v/\beta_1$: 12 cells) (***$p<10^{-3}$).

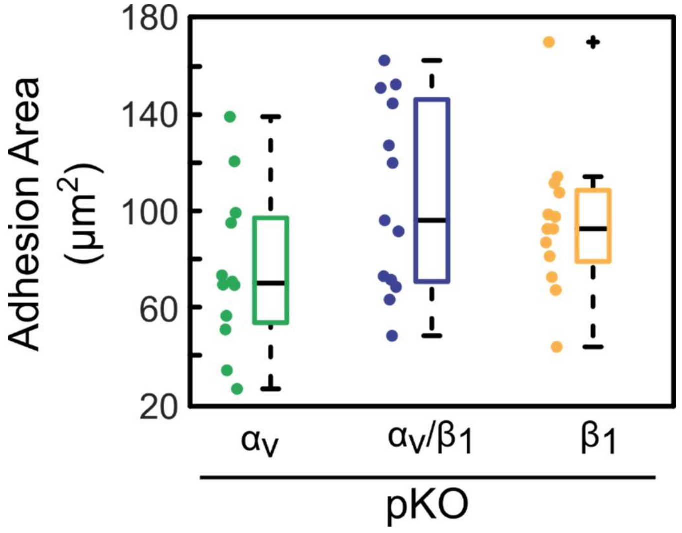


**Fig. S5. Adhesion area of pKO cells on MTS$_{FN9-10}$**

Adhesion area is calculated based on the thresholded GFP-paxillin signal. Differences in adhesion area were not significant between the three cell types, $p>0.01$.

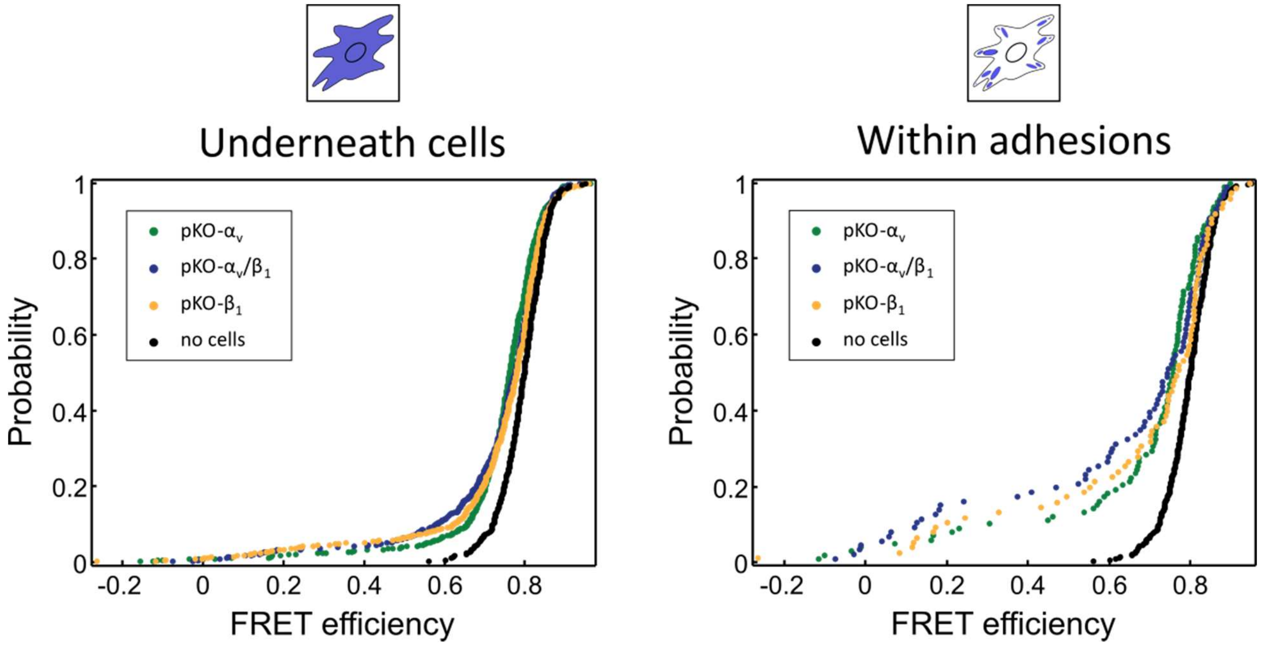


**Fig. S6. Single molecule FRET efficiency cumulative distributions pKO MEF cells adhering to $MTS_{FN9-10}$.**

Cumulative distributions of single-molecule FRET measurements for pKO-$\alpha_v$ (green), pKO-$\alpha_v$/$\beta_1$ (blue), pKO-$\beta_1$ (yellow) MEF cells, as well as no-load control (black) measurements, for all molecules underneath cells (left) and specifically within adhesions (right). pKO-$\alpha_v$: N = 106 cells, pKO-$\alpha_v$/$\beta_1$: N = 89 cells, pKO-$\beta_1$: 81 cells ; Underneath cells: pKO-$\alpha_v$: n = 714 molecules, pKO-$\alpha_v$/$\beta_1$: n = 610 molecules, pKO-$\beta_1$: 677 molecules; Within adhesions: pKO-$\alpha_v$: n = 98 molecules, pKO-$\alpha_v$/$\beta_1$: n = 86 molecules, pKO-$\beta_1$: 75 molecules, no-load: n = 299 molecules.

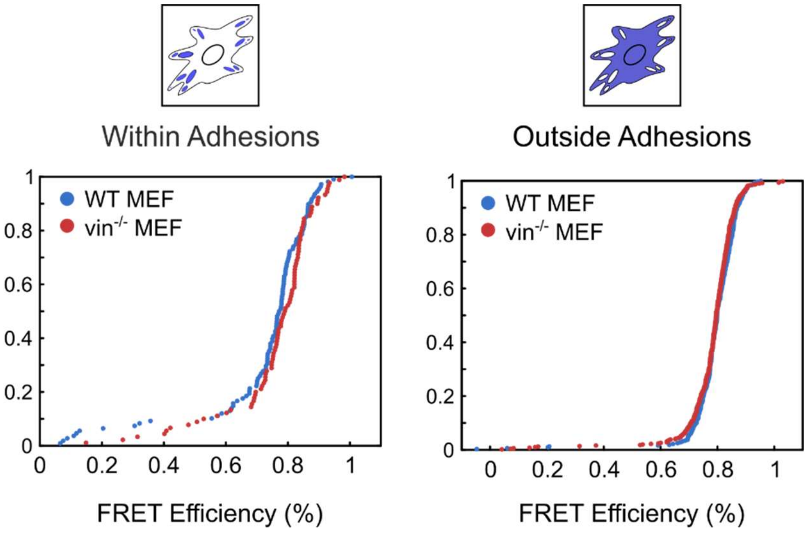


**Fig. S7. Single-molecule FRET efficiency CDFs for WT and vinculin KO MEF cells adhering to MTS$_{low}$.**

Cumulative distributions for single-molecule FRET measurements for WT and vinculin KO cells for: (A) within adhesions (WT: n = 108 molecules, N = 139 cells, vin$^{-/-}$: n = 90 molecules, N = 127 cells), and (B) outside of adhesions (WT: n = 320 molecules, N = 63 cells, vin$^{-/-}$: n = 500 molecules, N = 65 cells).

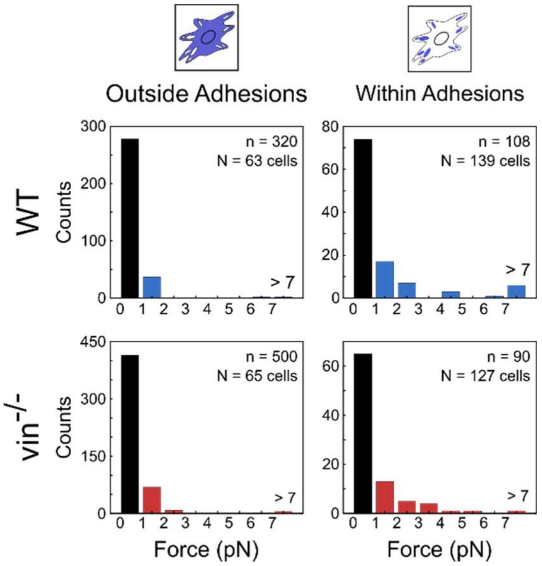


**Fig. S8. Single-molecule load distributions for WT and vin$^{-/-}$ MEFs adhering to MTS$_{low}$.** Histograms of the single-molecule load measurements for WT and vin$^{-/-}$ MEFs measured for cells adhering to MTS$_{low}$ for sensors outside adhesions (left) and within adhesions (right).

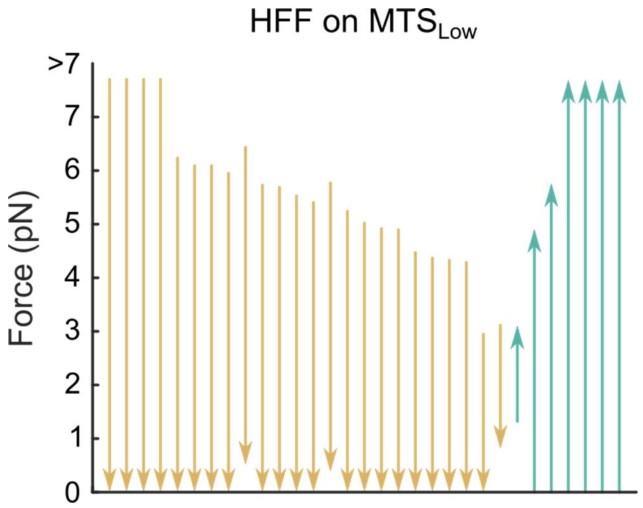


**Fig. S9. Dynamic step events for HFFs adhering to MTS$_{low}$.**

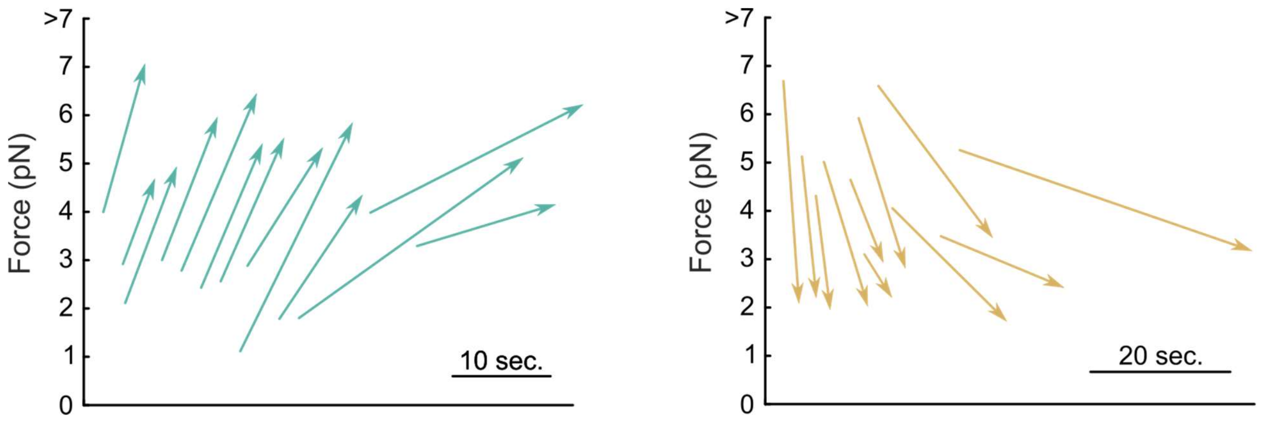


**Fig. S10. Dynamic ramp events for HFFs adhering to $MTS_{low}$.**

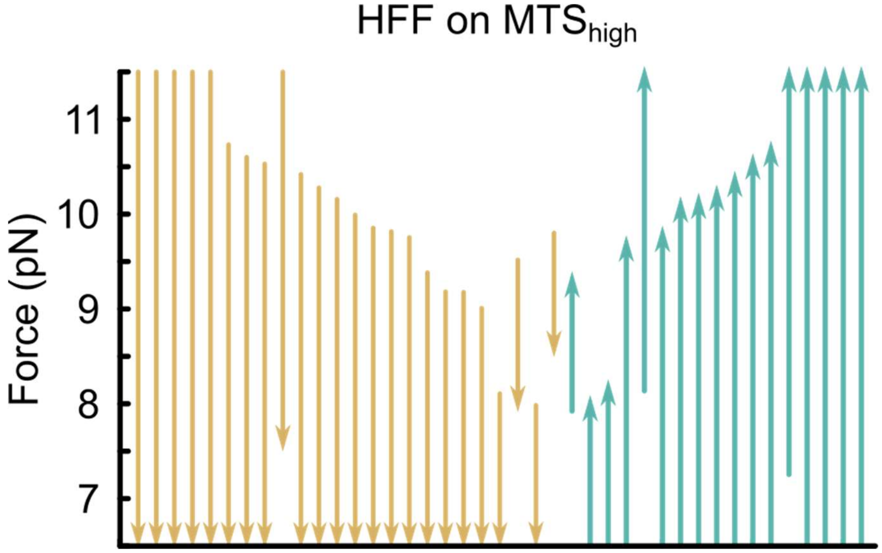


**Fig. S11. Dynamic step events for HFFs adhering to MTS$_{high}$.**

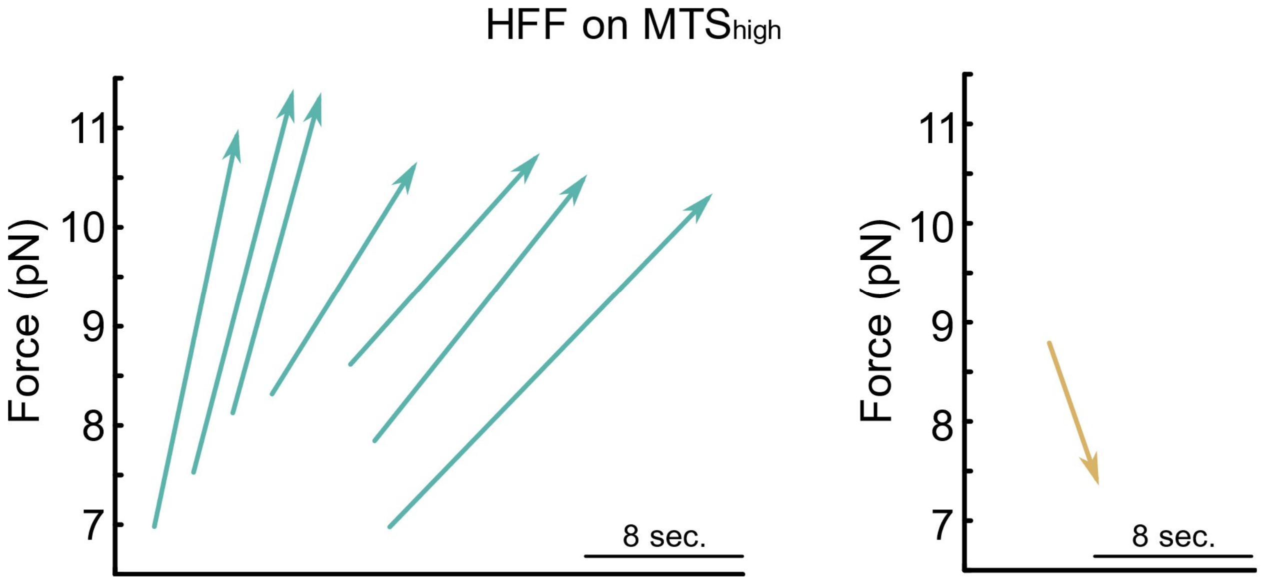


**Fig. S12. Dynamic ramp events for HFFs adhering to $MTS_{high}$.**

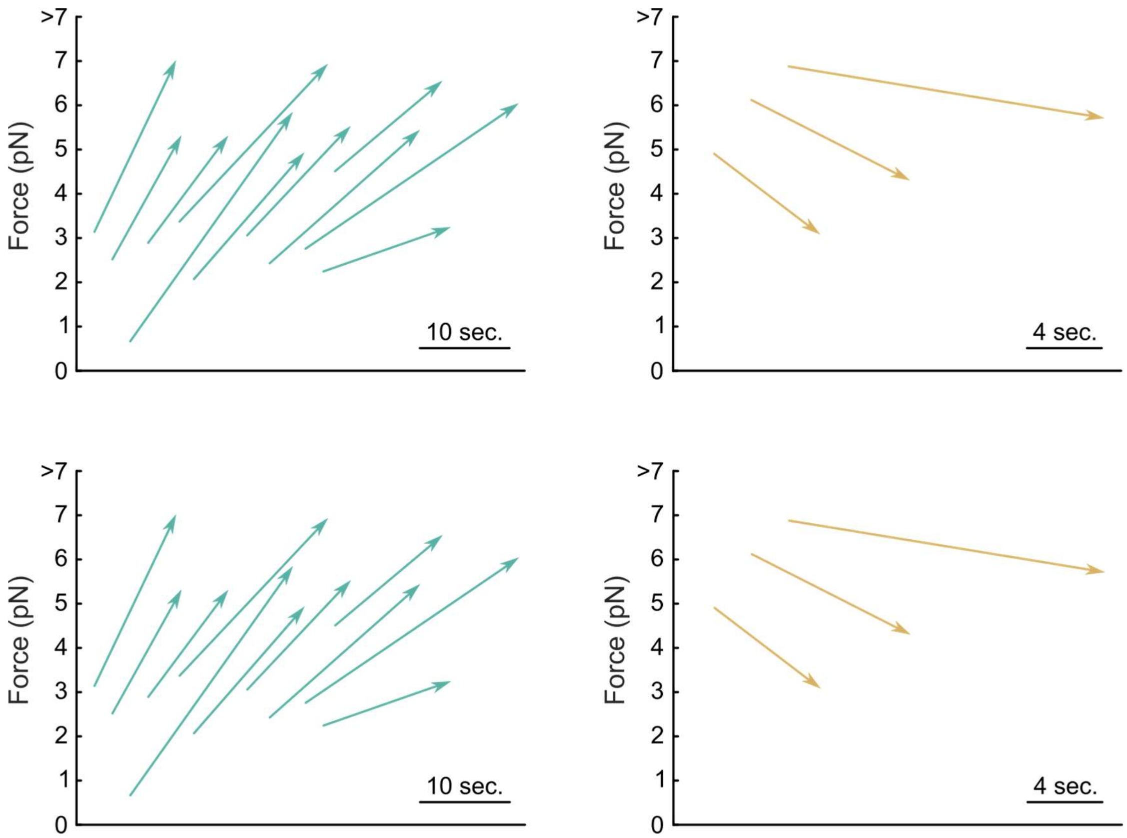


**Fig. S13. Dynamic ramp events for pKO-αv/β1 (top) and pKO-β1 (bottom) MEFs adhering to MTS$_{FN9-10}$.**

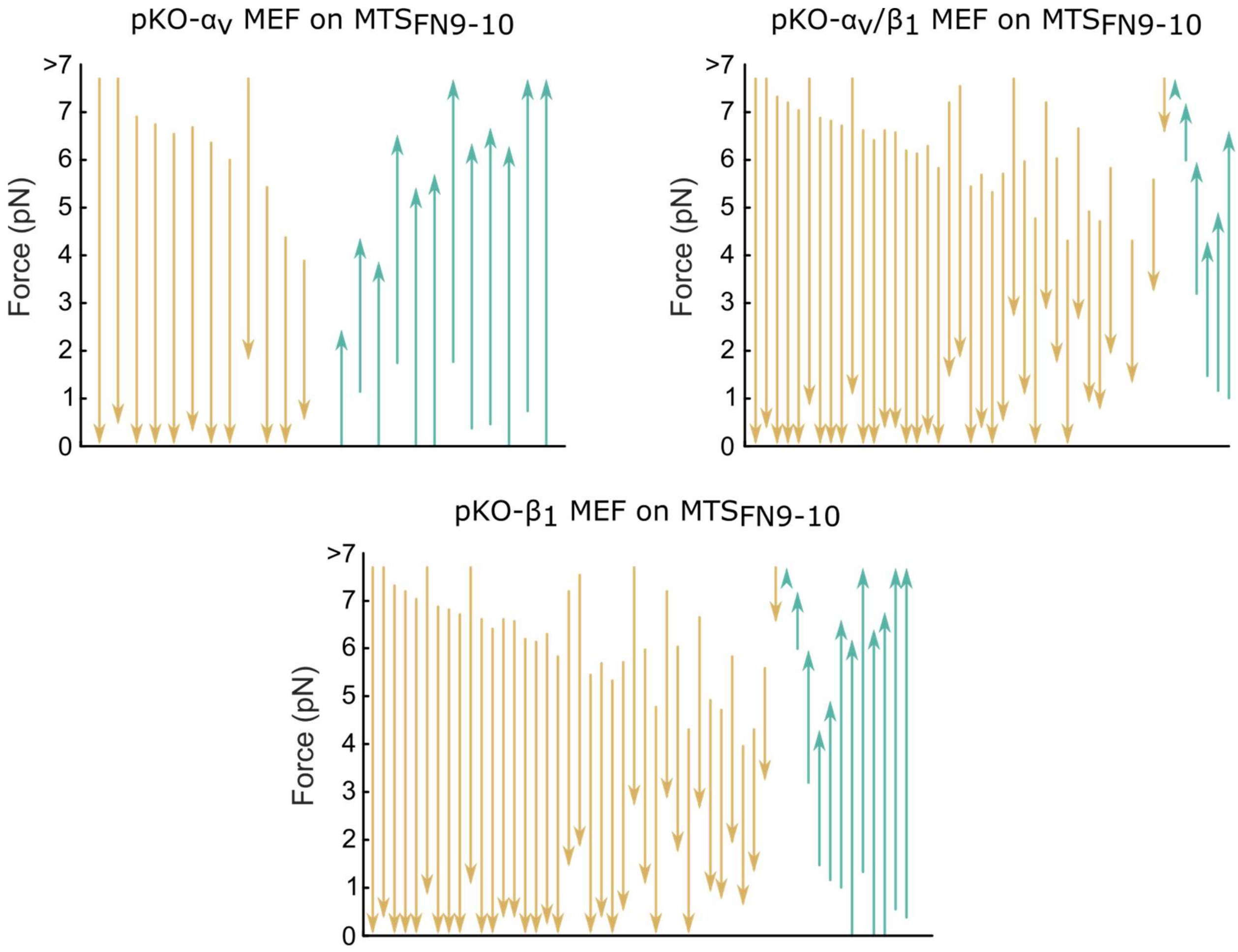


**Fig. S14. Dynamic step events for pKO MEF adhering to $MTS_{FN9\text{-}10}$.**

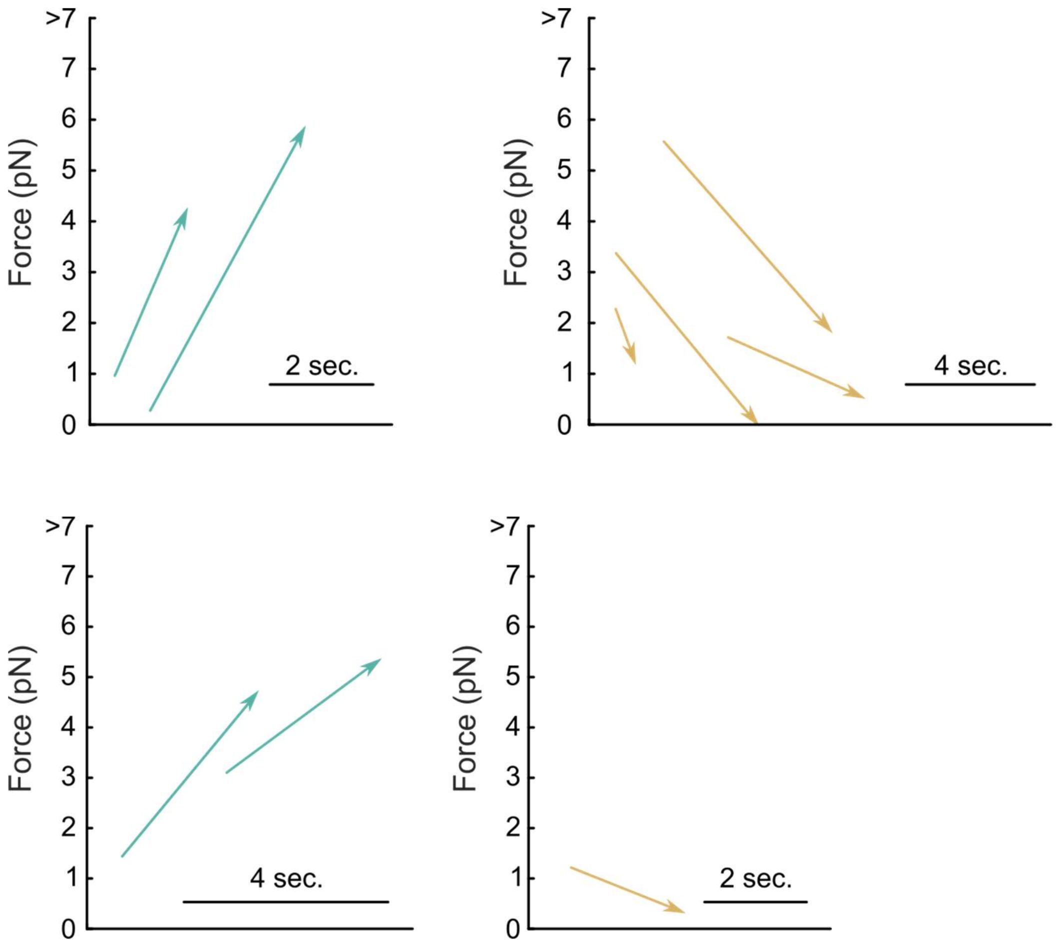


**Fig. S15. Dynamic ramp events for WT and vin$^{-/-}$ MEFs adhering to $MTS_{low}$.**

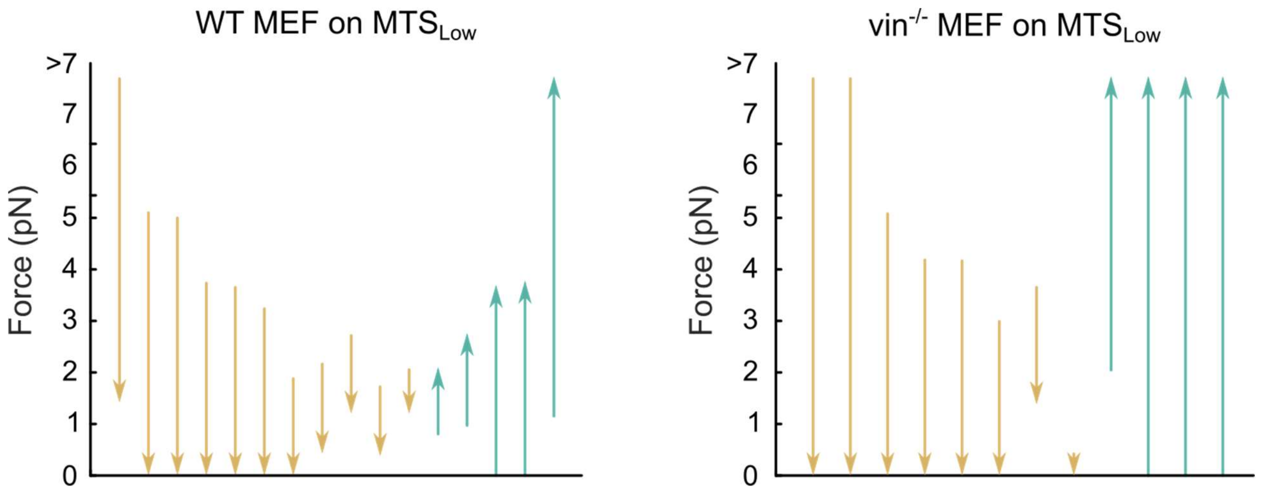


**Fig. S16. Dynamic step events for WT and vin$^{-/-}$ MEFs adhering to MTS$_{low}$.**

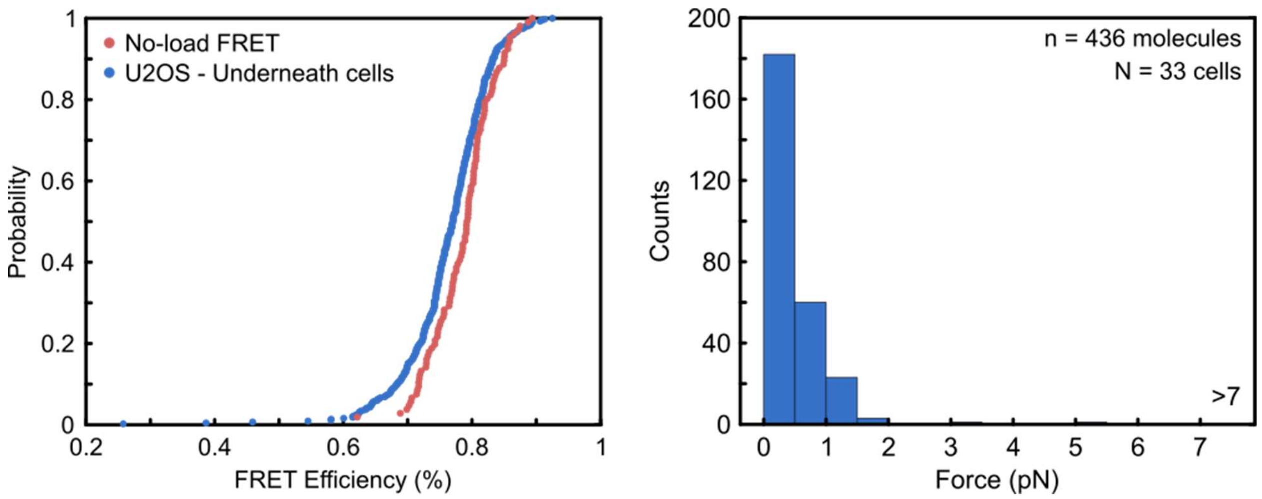


**Fig. S17. Single-molecule FRET efficiency CDF and force histogram for U2OS cells adhering to $MTS_{low}$.**

(A) Cumulative distribution for no-load sensors and sensors measured under U2OS cells. (B) Converted force values for sensors under U2OS cells.

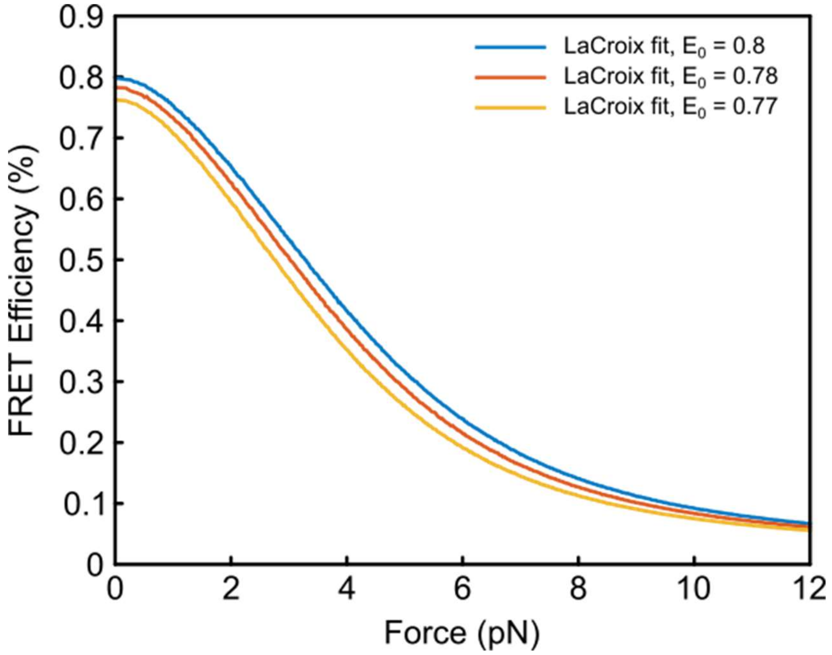


**Fig. S18. Updated FRET-force calibrations.**

FRET-force calibrations calculated for $GPGGA_8$-based tension sensors using updated calibrations calculated from LaCroix *et al*. (*47*). Different sensors and experimental conditions result in slightly different resting FRET efficiencies ($E_0$) and a new calibration curve is calculated to reflect each experimental condition.

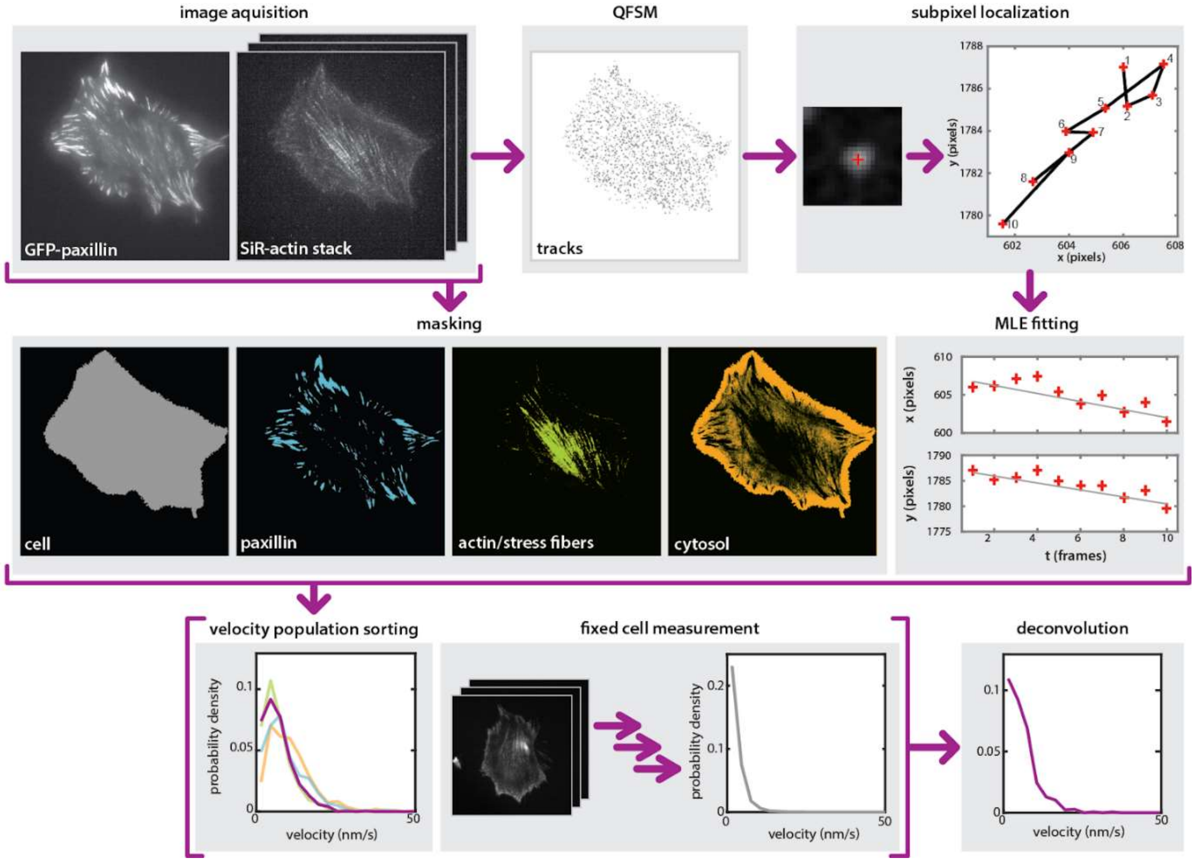


**Fig. S19. Actin tracking workflow.**

Images of GFP-paxillin expressing and SiR-actin labelled HFFs are acquired. A stack of SiR-actin images is fed into the quantitative fluorescent speckle microscopy (QFSM) software which identifies tracks with pixel-level precision. These tracks, together with the original images, are used to generate sub-pixel localized tracks, which are fit using maximum likelihood estimation (MLE) linear fits. The original images of both GFP-paxillin and SiR-actin are used to generate masks of cell edges, focal adhesions ("paxillin" mask), and stress fibers; and the "cytosol" mask is defined by regions within the cell that are excluded from the other two masks. The velocities from the MLE fitting are then sorted according to their origin point relative to the masks described above, yielding velocity histograms for each subcellular population. Using data from fixed cells (processed in the same way), the measurement noise is deconvolved from the velocity measurements to estimate the true underlying velocity distribution for each subpopulation.

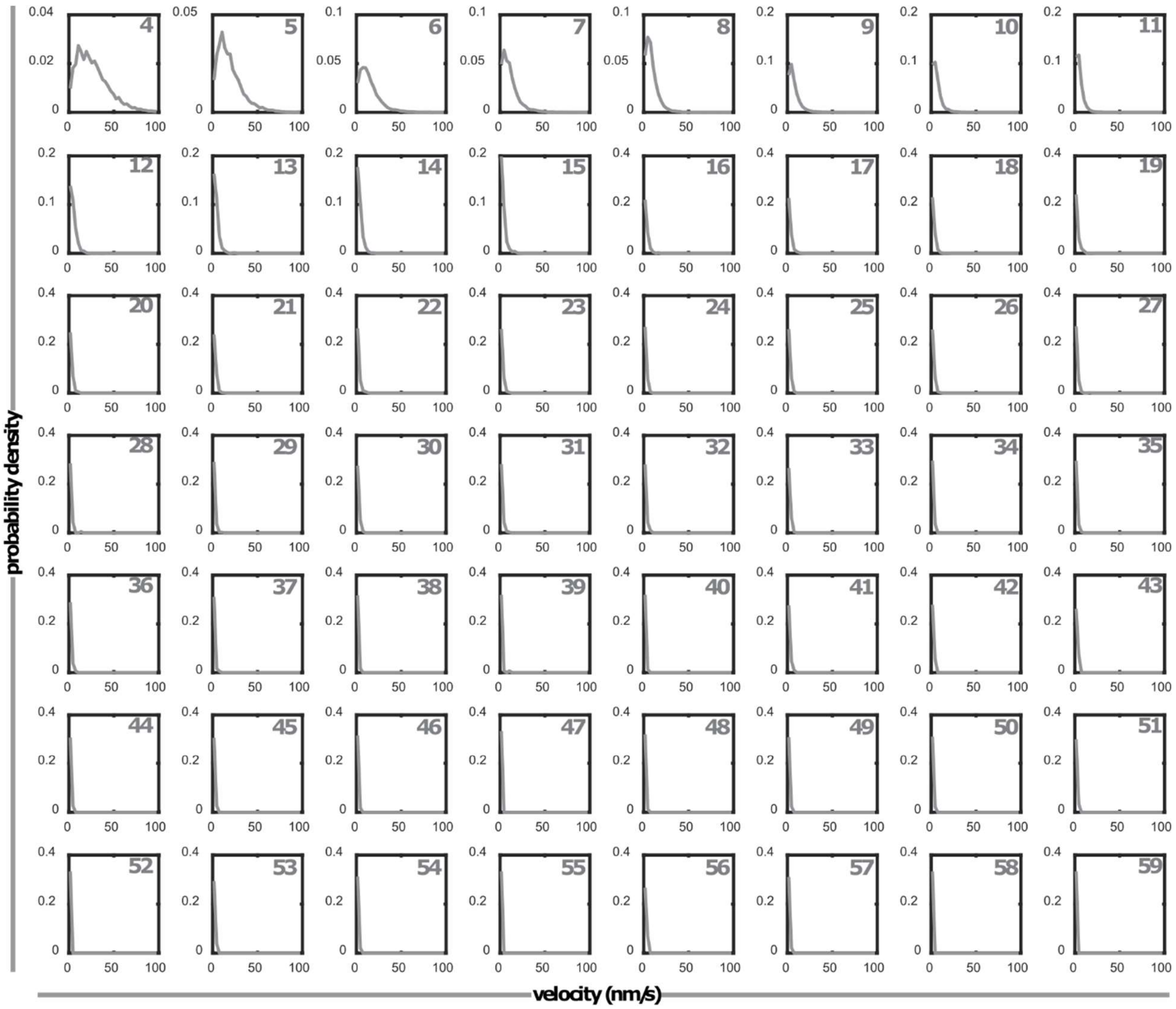


**Fig. S20. Velocity histograms for actin tracks in fixed cells, separated by track length.** The track length is indicated in the top right corner of each plot.

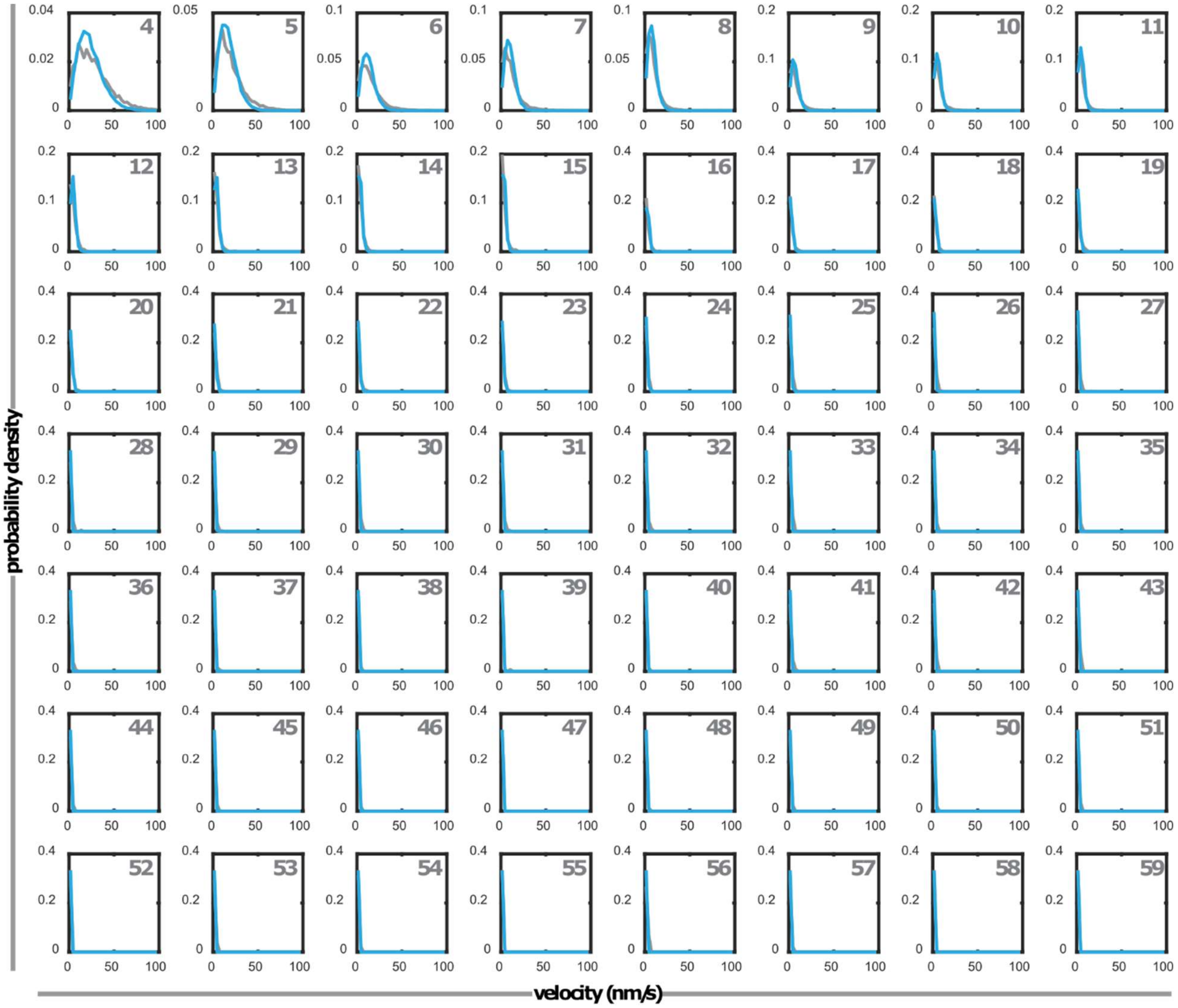


**Fig. S21. Simulations qualitatively capture the observed length-dependence of actin velocities for the control case (all true velocities equal to zero).**

Velocity histograms show the actin velocities in fixed cells (gray) and obtained by simulations (blue), separated by track length. Track lengths are indicated in the top right corner of each plot.

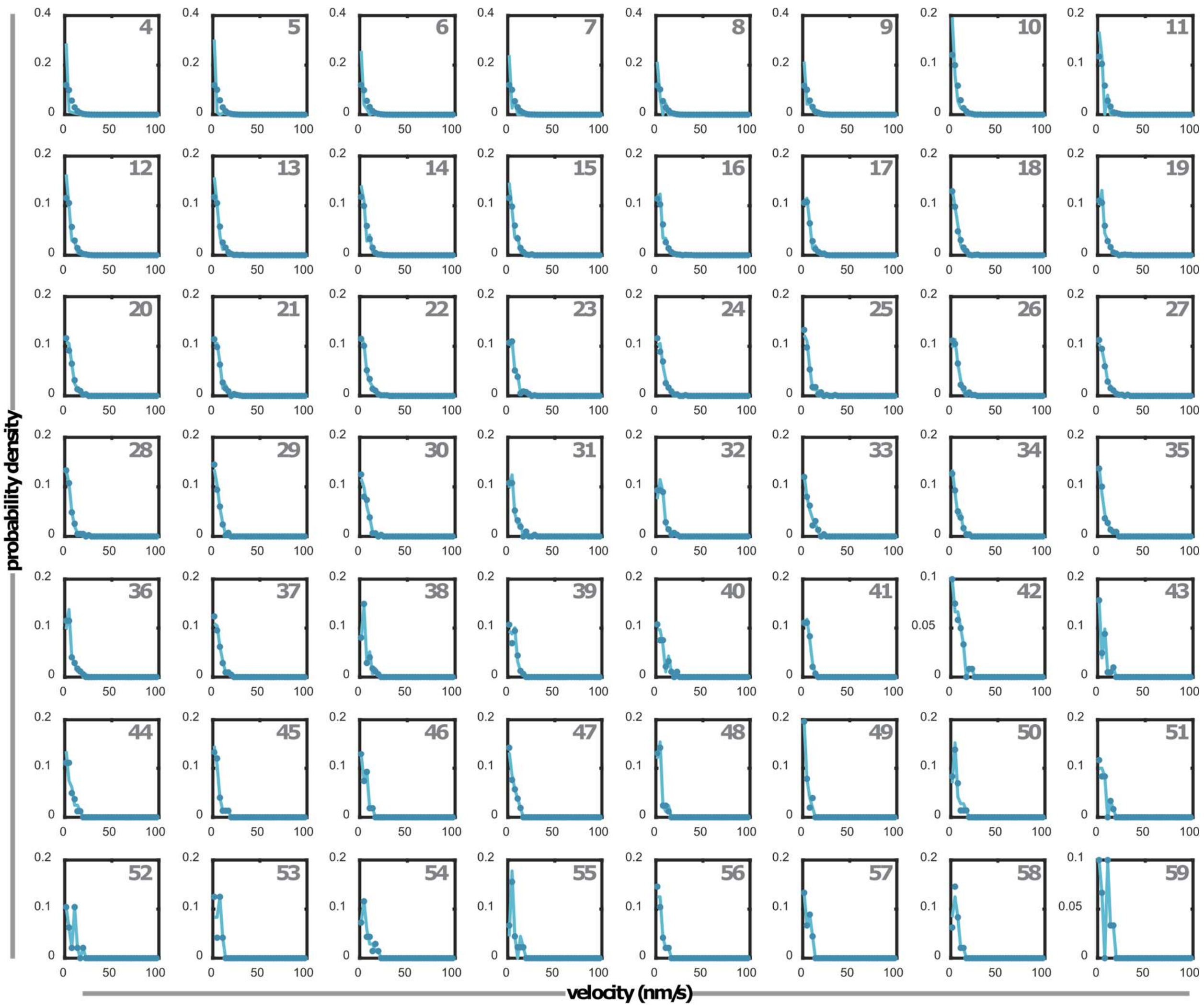


**Fig. S22. Deconvolution underestimates the velocity for short tracks.**

The true velocity distributions (without added noise; circles) from simulations and the resulting histograms deconvolved from the noisy velocity distributions (solid line). Velocities are separated by track length, indicated in the top right of each plot.

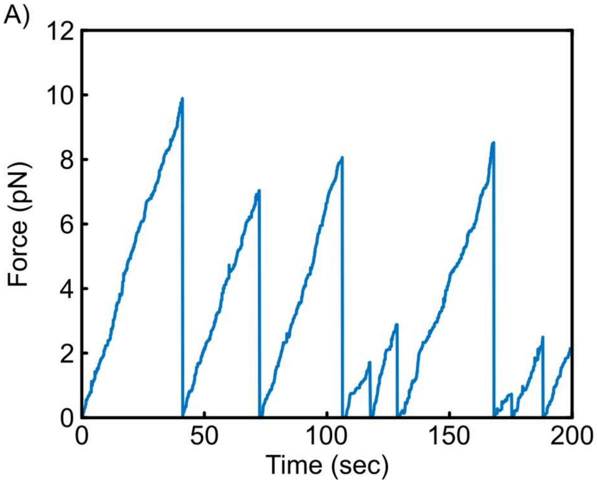


**Fig. S23. Clutch trace and force histogram for the base molecular clutch model.**
A) Force on a single clutch as predicted by the motor clutch model. B) Force distribution of bound clutches.

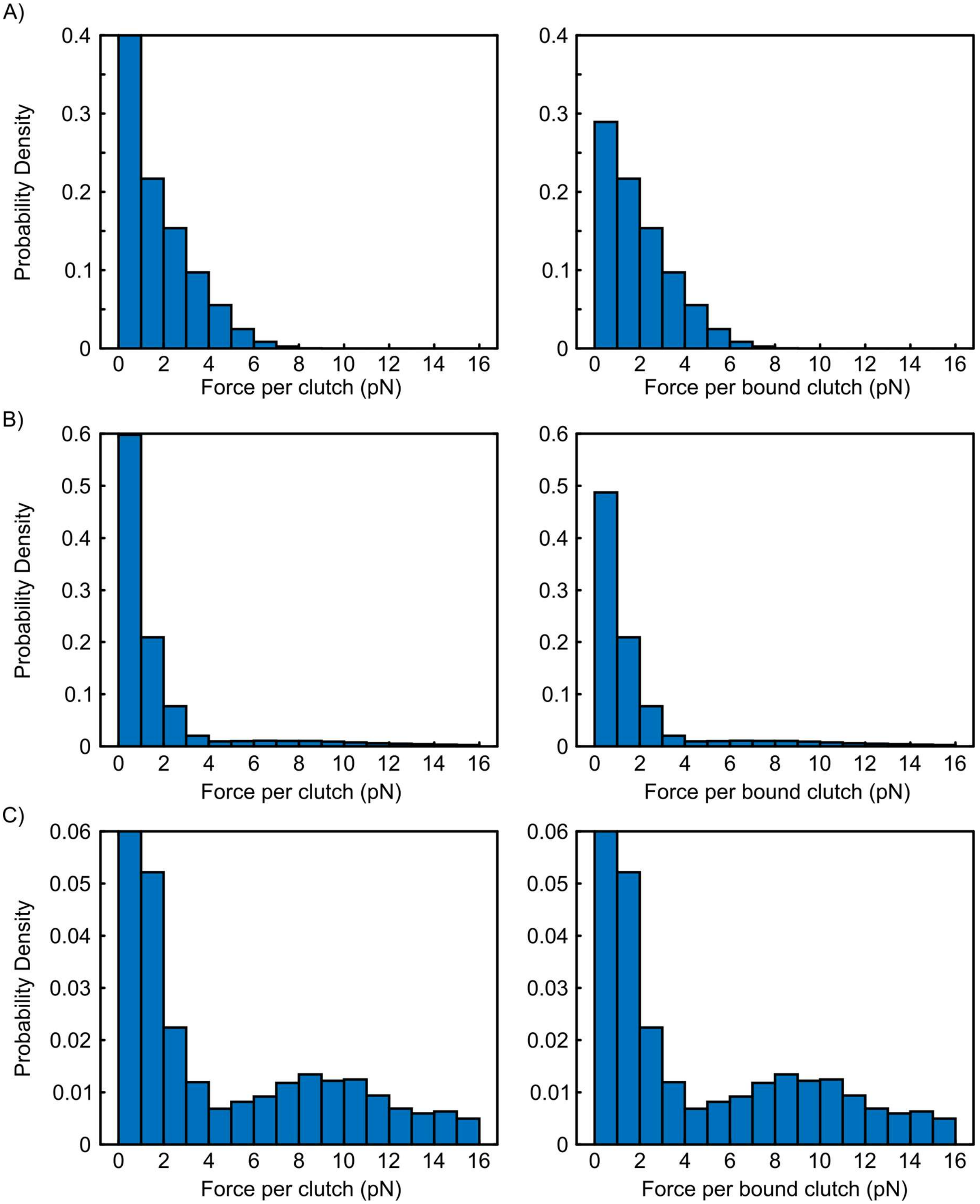


**Fig. S24. Force histograms for molecular clutch model simulations incorporating reinforcement or catch bond behavior.**

A) Force per clutch and force per bound clutch force distribution using a slip bond. B) Force per clutch and force per bound clutch force distribution using a slip bond with the inclusion of reinforcement leading to high force events. C) Force per clutch and force per bound clutch force distribution using a catch bond and reinforcement leading to a peak of high force events.

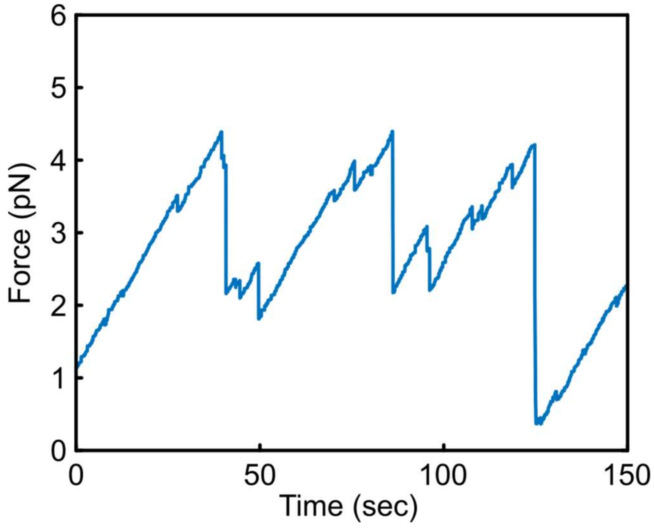


**Fig. S25. Time series for a single clutch experiencing viscous relaxation and rapid rebinding.**

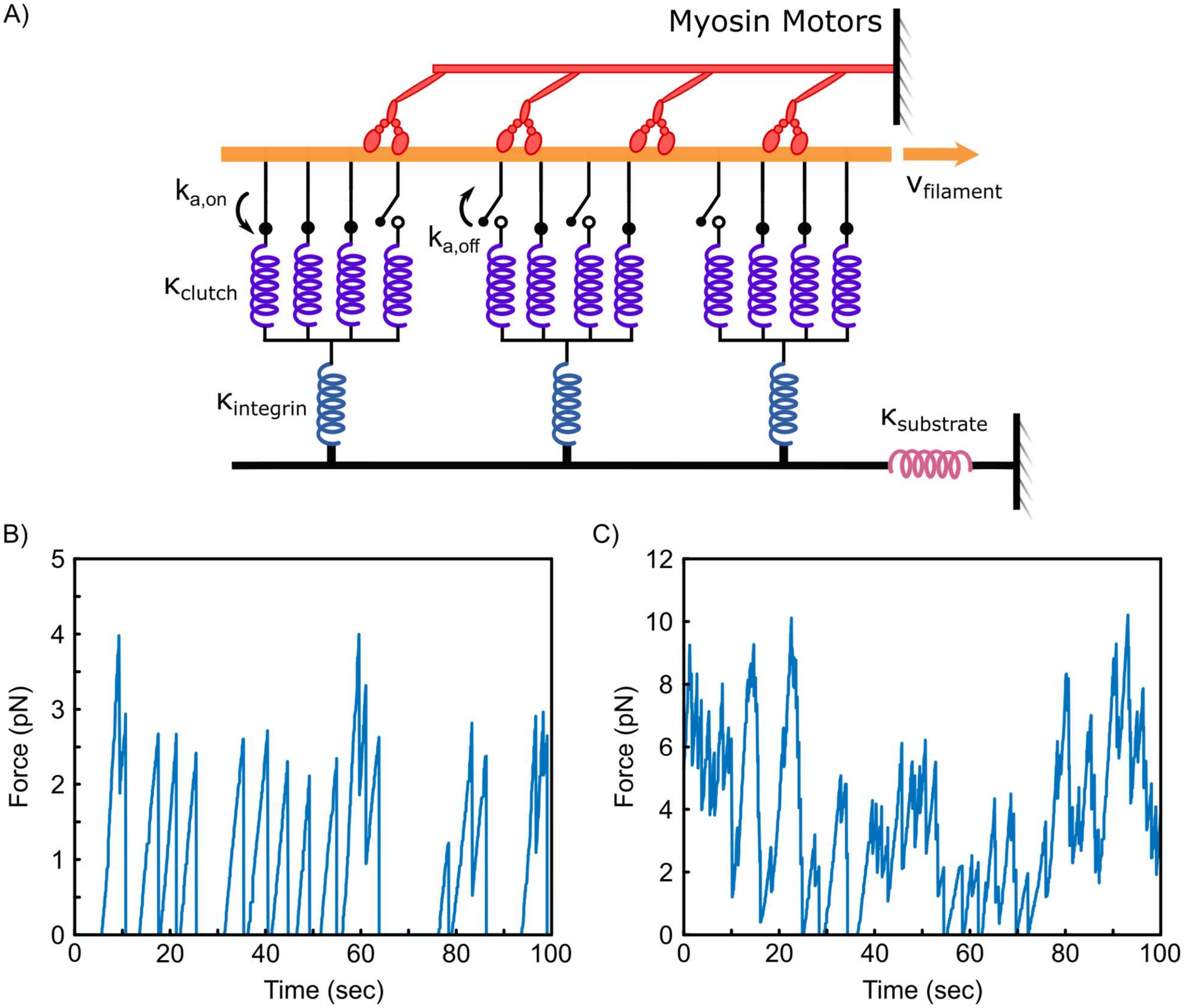


**Fig. S26. Molecular clutch model simulations incorporating multiple connections between integrins and actin.**

A) Schematic of motor clutch model including multiple connections between an integrin spring and actin. B) Time series for a single integrin when two clutches per integrin were used. C) Time series for a single integrin when twelve clutches per integrin were used.

| Cell type | # of unbound sensors (<2.5 pN) | # of bound sensors (>2.5 pN) | Significance Test ($\chi^2$ test) | p-value |
|---|---|---|---|---|
| pKO-$\alpha_v$ | 682 | 32 | pKO-$\alpha_v$ : pKO-$\alpha_v/\beta_1$ | ***7.3 x $10^{-6}$ |
| pKO-$\alpha_v/\beta_1$ | 543 | 67 | pKO-$\alpha_v$ : pKO-$\beta_1$ | ***7.3x$10^{-4}$ |
| pKO-$\beta_1$ | 616 | 61 | pKO-$\alpha_v/\beta_1$ : pKO-$\beta_1$ | (n.s.) 0.24 |

**Table S1. Number of bound and unbound sensors for pKO MEFs adhering to $MTS_{FN9-10}$ and calculated statistics.**

Populations of unbound vs. bound sensors were calculated as in Chang et al. (*14*). P-values were calculated using the $\chi^2$ significance test.

| | # of step unbinding events | # bound (>3pN or >7pN) | $F_{RI}$ | $k_I$ ($s^{-1}$) | $\tau_I$ (s) |
|---|---|---|---|---|---|
| HFFs on $MTS_{low}$ | 24 | 134 | 0.179 | 0.024 | 42 |
| HFFs on $MTS_{high}$ | 24 | 101 | 0.238 | 0.028 | 36 |
| pKO-$\alpha_v$ MEF on $MTS_{FN9\text{-}10}$ | 13 | 32 | 0.406 | 0.055 | 18 |
| pKO-$\beta_1$ MEF on $MTS_{FN9\text{-}10}$ | 39 | 61 | 0.639 | 0.144 | 7 |
| pKO-$\alpha_v/\beta_1$ MEF on $MTS_{FN9\text{-}10}$ | 24 | 67 | 0.358 | 0.045 | 22 |
| WT MEF on $MTS_{low}$ | 11 | 29 | 0.379 | 0.081 | 12 |
| $Vin^{-/-}$ MEF on $MTS_{low}$ | 8 | 37 | 0.216 | 0.036 | 27 |

**Table S2. Integrin-ligand lifetime estimates for different sensor-cell conditions.**

| Cell type | Sensor type | Steps (#events) | Average step increase (pN) | Average step decrease (pN) | Ramps (#events) | Average ramp increase (pN/sec) | Average ramp decrease (pN/sec) |
|---|---|---|---|---|---|---|---|
| HFF | $MTS_{low}$ | 31 | 6.1 (n = 7) | -5.4 (n = 24) | 24 | 0.39 (n = 13) | -0.61 (n = 11) |
| | $MTS_{high}$ | 41 | 8.2 (n = 17) | -8.3 (n = 24) | 8 | 0.49 (n = 7) | -1.1 (n = 1) |

**Table S3. Dynamic events observed for HFFs adhering to $MTS_{low}$.**

| Cell type | Sensor type | Steps (#events) | Average step increase (pN) | Average step decrease (pN) | Ramps (#events) | Average ramp increase (pN/sec) | Average ramp decrease (pN/sec) |
|---|---|---|---|---|---|---|---|
| pKO-$\alpha_v$ MEF | $MTS_{FN9-10}$ | 25 | 5.3 (n = 12) | -5.7 (n = 13) | 0 | -- | -- |
| pKO-$\beta_1$ MEF | | 51 | 4.6 (n = 12) | -5.3 (n = 39) | 15 | 0.29 (n = 10) | -0.39 (n = 5) |
| pKO-$\alpha_v/\beta_1$ MEF | | 34 | 5.7 (n = 10) | -5.6 (n = 24) | 14 | 0.23 (n = 11) | -0.20 (n = 3) |

**Table S4. Dynamic events observed for pKO MEFs adhering to $MTS_{FN9-10}$.**

| Cell type | Sensor type | Steps (#events) | Average step increase (pN) | Average step decrease (pN) | Ramps (#events) | Average ramp increase (pN/sec) | Average ramp decrease (pN/sec) |
|---|---|---|---|---|---|---|---|
| WT MEF | $MTS_{low}$ | 16 | 3.3 (n = 5) | -3.0 (n = 11) | 6 | 2.1 (n = 2) | -0.88 (n = 4) |
| $vin^{-/-}$ MEF | | 12 | 7.0 (n = 4) | -4.3 (n = 8) | 3 | 0.99 (n = 2) | -0.4 (n = 1) |

**Table S5. Dynamic events observed for WT and $vin^{-/-}$ MEFs adhering to $MTS_{low}$.**

| Sensor Type | Total # of sensors | # of dynamic-like sensors |
|---|---|---|
| $MTS_{low}$ | 550 | 4 |
| $MTS_{high}$ | 260 | 2 |
| $MTS_{FN9-10}$ | 224 | 2 |

**Table S6. Number of dynamic-like sensors from no-load measurements.**

| **Cell type** | **Sensor type** | **Total # sensors** | **# sensors Ramp loading** | **# sensors Ramp unloading** | **# sensors Step loading** | **# sensors Step unloading** |
|---|---|---|---|---|---|---|
| HFF | $MTS_{low}$ | 781 | 11 | 9 | 5 | 22 |
| HFF | $MTS_{high}$ | 595 | 7 | 1 | 9 | 11 |
| pKO-$\alpha_v$ MEF | $MTS_{FN9-10}$ | 714 | - | - | 9 | 11 |
| pKO-$\beta_1$ MEF | $MTS_{FN9-10}$ | 677 | 9 | 3 | 9 | 23 |
| pKO-$\alpha_v$/$\beta_1$ MEF | $MTS_{FN9-10}$ | 610 | 7 | 3 | 12 | 36 |
| WT MEF | $MTS_{low}$ | 428 | 1 | 1 | 3 | 8 |
| $Vin^{-/-}$ MEF | $MTS_{low}$ | 590 | 1 | 4 | 5 | 11 |

**Table S7. Number of dynamic sensors.**

| | |
|---|---|
| $x_{cl,t}$ | Clutch displacement |
| t | time |
| $K_c$ | Clutch spring constant |
| N | Damping constant |
| C | Rebinding constant |
| $k_{on}$ | Clutch on-rate constant |

**Table S8. Parameter definitions in the viscous relaxation and rapid rebinding model.**

| Parameter | Symbol | Value | Units | Reference(s) |
|---|---|---|---|---|
| Number of motors | $n_m$ | 200 | - | (*12*) |
| Motor unloaded velocity | $V_u$ | -120 | nm/s | (*12*) |
| Motor force | $F_m$ | 2 | pN | (*12*) |
| Number of clutches | $N_c$ | 120 | - | (*12*) |
| Clutch on-rate | $k_{a,on}$ | 1 | 1/s | (*12*) |
| Basal clutch off-rate | $k_{a,off}$ | 0.03 | 1/s | (*12*) |
| Characteristic bond rupture force | $F_b$ | 8 | pN | (*12*) |
| Clutch stiffness | $K_c$ | 0.16 | pN/nm | (*12*) |
| Substrate stiffness | $K_s$ | $10^6$ | pN/nm | - |
| Crosslinker unbinding rate (irreversible) | $k_{x,on}$ | 0 | 1/s | (*12*) |
| Crosslinker unbinding rate (reversible) | $k_{x,on}$ | 10 | 1/s | (*49*) |
| Crosslinker binding rate | $k_{x,off}$ | 100 | 1/s | (*49*) |

**Table S9. Parameters for the updated clutch model including reversible crosslinkers.**
